# Structure-function dissection of huntingtin exon 1 identifies a PRD-driven modifier of neuronal toxicity in Huntington’s disease

**DOI:** 10.64898/2026.07.23.740298

**Authors:** Raffaele Iennaco, Camilla Maffezzini, Simone Maestri, Andrea Scolz, Angela Cattaneo, Alessio Murgia, Camilla Trovesi, Eugenia Cammarota, Elena Vezzoli, Andrea Falqui, Dan P. Felsenfeld, Thomas F. Vogt, Angela Bachi, Chiara Zuccato, Elena Cattaneo

## Abstract

The *Huntingtin* gene (*HTT*) contains a conserved, yet expandable CAG repeat within exon 1. While the pathogenic expansion in Huntington’s Disease (HD) is well studied, the role of surrounding domains remains unclear. Using genome-edited mini-organoids and neurons, we dissected *HTT* exon 1 and found species-specific toxicity: the human variant caused more severe deficits than the mouse. Swapping the proline-rich domain (PRD) - the most divergent region - revealed its key role: the mouse PRD mitigated, while the human PRD worsened neuronal phenotypes. Omics profiling showed that pathogenic human exon 1 induced broad protein dysregulation, largely reversed by mouse PRD replacement. Bioinformatics implicated the actin cytoskeleton and transcriptional coactivator MKL2/MRTFB. We validated MKL2/MRTFB dysregulation in HD models and showed that restoring its expression rescued neuronal abnormalities. These findings highlight the PRD’s contribution to HD toxicity and point to MKL2/MRTFB and the cytoskeleton as candidate mediators.

## Introduction

The huntingtin protein (HTT) contains a poly-glutamine (polyQ) tract that, in humans, is typically encoded by a stretch of pure CAG repeats followed by a CAACAG sequence within exon 1 of the *HTT* gene. This polyQ region has been extensively studied, as its abnormal expansion beyond 36 repeats causes Huntington’s Disease (HD), a progressive neurodegenerative disorder^1^. In humans, the polyQ tract is flanked by two highly conserved domains within exon 1: a 17-amino acids N-terminal segment (N17) and a downstream proline-rich domain (PRD)^2,3^. The PRD begins with a polyproline (polyP) tract (P1), encoded by *(i)* an intervening CCGCCA sequence, *(ii)* a stretch of pure CCG repeats, and *(iii)* a few CCT repeats. A second uninterrupted polyP tract (P2) follows within the broader PRD^4^. This repetitive domain ends with a short 10-amino-acid C-terminal segment (C-term), completing exon 1.

While polyQ length is a major determinant of HD onset and severity, it does not fully explain the wide inter-individual variability in age of onset and disease severity^5,6^, suggesting the influence of additional genetic factors both within and outside the polyQ-encoding region^7–9^. Genome-wide association studies (GWASs) have since identified numerous *cis-* and *trans*-modifiers of HD pathogenesis^10^, including those associated with somatic CAG repeat instability and DNA maintenance pathways^11–15^. These findings support a two-step model of HD pathogenesis: an initial phase of somatic expansion, followed by neuronal degeneration driven by the resulting toxicity^16^.

However, other human genetic studies have identified *HTT* variants near the CAG repeat that accelerate disease onset without increasing somatic instability^17,18^, indicating that additional, somatic instability-independent mechanisms also contribute to HD pathogenesis. Among these, the loss of the intervening CCGCCA motif within the P1 tract of the PRD has been associated with earlier onset^17^, refocusing attention on the PRD as a potential modulator of HTT toxicity. Supporting this view, the PRD has been implicated in aggregation, fibril formation, and HTT solubility^4,19–21^. Experimental interventions targeting the PRD – such as PRD deletions or sequence-specific antibodies – have shown reduced toxicity, likely by enhancing mutant HTT turnover and lowering its intracellular levels^22,23^. In-cell analyses and human studies have revealed a beneficial role for longer CAG (Q) repeats in the *HTT* gene, lending support to the theory of antagonist pleiotropy in HTT activity^24,25^. Evolutionary studies further suggest that the emergence of the PRD and the expansion of the polyQ tract occurred concomitantly during vertebrate evolution^24^, pointing to a possible co-evolution and functional interdependence between these two domains. Moreover, polymorphisms in the uninterrupted CCG repeats adjacent to the polyQ tract have been associated with CAG repeat variability in both control and HD populations^26,27^.

Altogether, the PRD emerges as a key *cis*-modifier in HD pathogenesis, potentially influencing both CAG instability and HTT-related toxicity at the DNA, RNA and protein levels.

Despite growing interest in *cis*-modifiers, the specific contributions of exon 1 domains – beyond the polyQ tract – to quantifiable HD phenotypes remain largely unexplored. Moreover, *in vitro* systems enabling domain-specific dissection of HTT toxicity are lacking. Here, we leveraged a Recombination-Mediated Cassette Exchange (RMCE)-based ES cell system^24^, in which the endogenous mouse huntingtin *(Htt)* exon 1 *locus* is replaced by an RMCE element, to generate a panel of knock-in cell lines expressing distinct, unmodified versions of *HTT* exon 1 (named “HuntEx1” platform). Using this platform, we investigated the differential pathogenic potential of human versus mouse exon 1 sequences, which primarily diverge in the PRD, and the contribution of the PRD to HD-related neuronal phenotypes. We reasoned that identifying sequence-specific effects on cellular pathology and molecular dysregulation could reveal novel modulators of HD severity and inform therapeutic strategies.

## Results

### Mouse and human HTT exon 1 differentially affect neurogenesis

HD mouse models expressing a chimeric huntingtin protein with human exon 1 exhibit earlier pathology than those expressing fully murine Htt^28^, suggesting increased toxicity of the human sequence. To investigate this in a controlled genetic background, we used a genome-edited, mouse ES cell system enabling parallel assessment of distinct *HTT* exon 1 variants. Specifically, we generated knock-in lines expressing either the mouse or human *HTT* exon 1 with non-pathogenic or pathogenic polyQ lengths, integrated at the endogenous murine *Htt locus* on one allele, while the second allele remained unmodified. This was achieved using a previously established RMCE-based platform^24^, which allows site-specific integration at the exon 1 *locus*. The resulting isogenic lines expressed murine exon 1 with 7Q (non-pathogenic, mWT/+), or 107Q (pathogenic, mHD/+), and human exon 1 with 20Q (non-pathogenic, hWT/+) or 72Q (pathogenic, hHD/+) (**Fig. 1a**). Lines carrying human exon 1 at the murine *locus* are referred to as “humanized”. Correct flp-mediated RMCE cassette exchange was confirmed by PCR and sequencing of exon 1, while HTT expression was evaluated by western blot analysis (**Extended Data Fig. 1a**). Maintenance of pluripotency after editing was validated by immunofluorescence for OCT3/4 and SOX2 (**Extended Data Fig. 1b**).

**Fig. 1:**
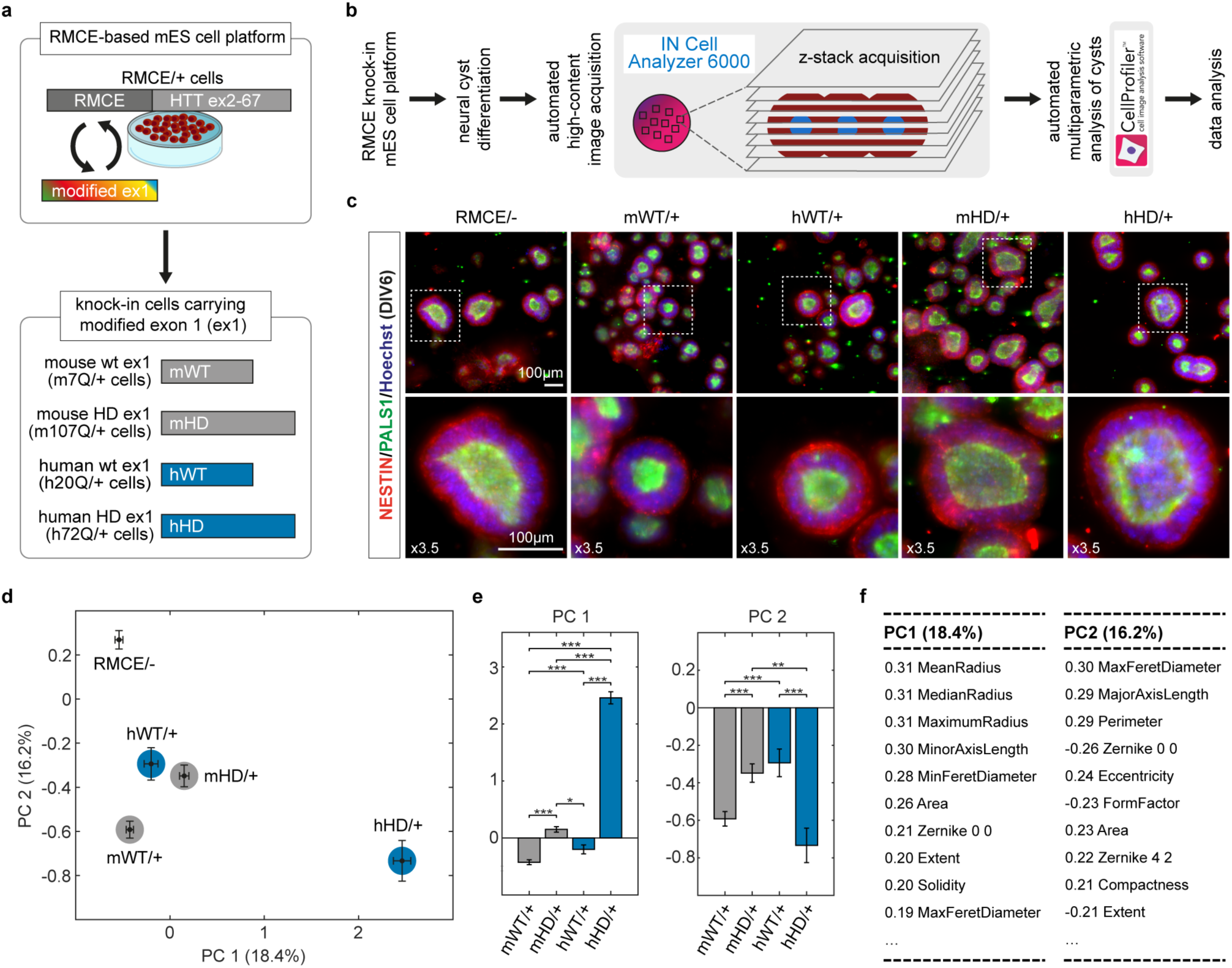
Species-specific impact of *HTT* exon 1 in the neural cyst assay. **a**, Schematic representation of RMCE-based generation of knock-in mESC lines carrying mouse or human *HTT* exon 1 with either wild-type or pathological polyQ lengths. **b**, Workflow of the automated multiparametric analysis pipeline used in the neural cyst assay. **c**, Representative images (*top row*) of neural cysts at DIV6 derived from RMCE/-, mWT/+, hWT/+, mHD/+, and hHD/+ mESC lines, stained for NESTIN (red), PALS1 (green), and Hoechst (blue). Scale bar:100 μm. Insets (*bottom row*) are shown at 3.5X magnification. **d**, Principal component analysis (PCA) of multiparametric cyst phenotypes, showing PC1 (18.4%) and PC2 (16.2%) variance explained. **e**, Bar plots displaying PC1 and PC2 values. Each dot (**d**) and bar (**d**,**e**) represent the median ± SEM from n>3 independent experiments. Statistical analysis by Kruskal-Wallis test with Bonferroni correction: ** P<0.01, *** P<0.001. **f**, Ranked contribution of individual neural cyst parameters to PC1 and PC2.

Given HTT’s physiological role in neural development^2,3^, and recent evidence linking HD mutation to early neurodevelopmental defects^29–34^, we tested whether these exon 1 variants differentially affect neural differentiation. We used a 3D neural cyst system – polarized neuroepithelial organoids resembling the developing neural tube – to model early neurogenesis. After 6 days in single-cell suspension (adapted from Meinhardt *et al.*, 2024^35^), all lines formed neural cysts with characteristic radial architecture, including apico-basal polarity and microvilli lining the single central lumen, as confirmed by fluorescence and electron microscopy (**Extended Data Fig. 2a-c**). To analyze cyst morphology, we employed high-content automated image acquisition coupled with multiparametric quantification (**Fig. 1b** and **Extended Data Fig. 2d**). Immunostaining for NESTIN, a neural progenitor marker, and PALS1, an apical polarity protein, confirmed the formation of properly polarized neurepithelial cysts (**Fig. 1c**). Principal component analysis (PCA) of size and shape parameters revealed that *Htt* knock-out cysts (RMCE/-) were morphologically distinct from wild-type murine exon 1 cysts (mWT/+) (**Fig. 1d**), reinforcing the essential role of huntingtin in early neurogenesis and confirming previous findings showing that loss of wild-type Htt disrupts radial neural structures^24,36^. Importantly, cysts carrying the murine HD exon 1 (mHD/+) diverged from mWT/+ controls along both principal components (PC1: size; PC2: shape) (**Fig. 1d-f**). Cysts expressing wild-type human exon 1 (hWT/+) also differed from mWT/+ cysts, suggesting that the human exon 1 sequence – even with a non-pathogenic polyQ length – can alter early neural morphology. Importantly, cysts expressing pathogenic exon 1 (mHD/+ and hHD/+) displayed distinct morphological profiles not only compared to their respective wild-type controls but also from each other, indicating that mouse and human pathogenic exon 1 variants have differential effects on phenotype (**Fig. 1d,e)**. These results point to a greater detrimental potential of human HD exon 1 compared to the murine version. Finally, both HD cyst populations were morphologically distinct from *Htt* knock-out cysts (**Fig. 1d**), supporting a gain-of-function mechanism for mutant huntingtin at this developmental stage.

### The PRD confers species-specific toxicity to human HD exon 1

To identify the domain responsible for this differential toxicity, we aligned and compared the amino acid (aa) sequences of murine and human exon 1. The N17 region is conserved (17 aa in both), with only minor synonymous substitutions (**Fig. 2a**). The polyQ region differs in length and codon usage: murine sequences consistently contain 7 Q with a CAA codon in the third position, whereas human wild-type sequences vary in length and typically include a penultimate CAA interruption (**Fig. 2a**). The most pronounced divergence lies in the PRD: it consists of 34 aa in mouse and 38 aa in human, with distinct numbers, positions and lengths of uninterrupted polyproline stretches. Specifically, the murine PRD contains four uninterrupted polyproline repeats separated by non-proline residues, while the human PRD includes only three such tracts, located in different positions and composed of distinct repeat lengths (**Fig. 2a**). The interspersed amino acid sequences between proline stretches also differ between the two species. Finally, at the 3’-end of exon 1, the C-terminal region comprises 8 aa in mouse and 10 in human, with the additional residues in the human sequence being valine (V) and alanine (A). The human C-terminal coding sequence also contains three synonymous substitutions relative to the murine one (**Fig. 2a**).

**Fig. 2:**
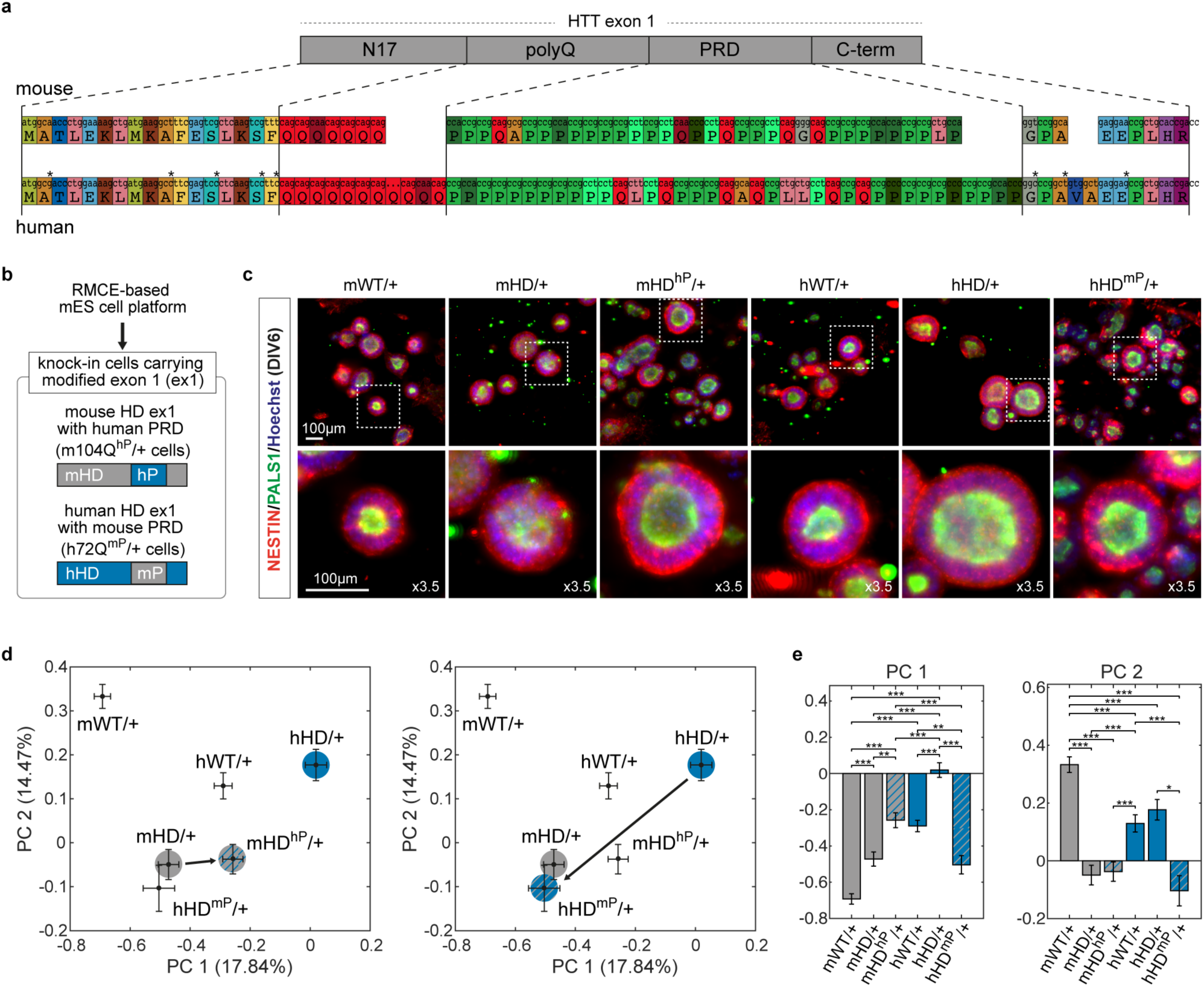
PRD swapping modifies the impact of HTT exon 1 on neural cysts formation. **a**, Comparative alignment of mouse and human *HTT* exon 1 sequences at the nucleotide and amino acid levels. The mouse sequence is based on GRCm39 (Genome Reference Consortium Mouse Build 39), and the human sequence on GRCh38 (Genome Reference Consortium Human Build 38). Different colors indicate distinct nucleotide triplets or amino acids, asterisks denote synonymous substitutions. **b**, Schematic of the generation of knock-in mESC lines carrying chimeric HTT exon1 constructs: mHD^hP^ (mouse HD exon 1 with human PRD), and hHD^mP^ (human HD exon 1 with mouse PRD. **c**, Representative images (*top row*) of neural cysts derived from mWT/+, mHD/+, mHD^hP^/+, hWT/+, hHD/+, and hHD^mP^/+ mESC lines stained for NESTIN (red), PALS1 (green), and Hoechst (blue) at DIV6. Scale bar: 100 μm. Insets (*bottom row*) show 3.5X magnification. **d**, PCA of multiparametric neural cyst phenotypes, showing PC1 (17.84% of variance) and PC2 (14.47%). Left panel: mHD^hP^/+ cysts shift away from mHD/+ toward hHD/+ profiles. Right panel: hHD^mP^/+ cysts diverge from hHD/+ and cluster to mHD/+ cysts. **e**, Bar plots showing PC1 and PC2 values. Each dot (**d**) and bar (**d**,**e**) represent the median ± SEM from n>3 independent experiments. Kruskal-Wallis test with Bonferroni correction: * P<0.05, ** P<0.01, *** P<0.001. Contributions of individual neural cyst parameters to PC1 and PC2 are shown in **Extended Data** Fig. 2e.

This divergence led us to hypothesize that the PRD is a key driver of species-specific HTT toxicity. To test this, we engineered chimeric knock-in mES cell lines by swapping the PRD between species. We inserted the human PRD into the murine HD exon 1 context (mHD^hP^/+) and, conversely, replaced the human PRD with the murine one (hHD^mP^/+), generating reciprocal constructs to rigorously test its functional impact (**Fig. 2b**). All lines were validated for correct RMCE cassette exchange, HTT expression and pluripotency (**Extended Data Fig. 1**), before neural cyst induction (**Fig. 2c**). Morphological profiling showed that mHD^hP^/+ cysts differed significantly from mHD/+, indicating that inserting the human PRD into a mouse background increases pathogenicity (**Fig. 2d,e**; *left panels*). Conversely, hHD^mP^/+ cysts were less morphologically abnormal than hHD/+ cysts, with improvements in both size and shape parameters (**Fig. 2d,e**; *right panels*), indicating that the murine PRD mitigates human HD exon 1 toxicity.

### The PRD affects dendritic architecture and neuronal function in HD neurons

We next asked whether these effects extend to mature neuronal phenotypes. All cell lines were differentiated into neurons using an established protocol^37^ (**Fig. 3a**), yielding a predominantly dorsal cortical identity, as indicated by the expression of TBR1, CTIP2, and VGLUT2, and the absence of the ventral marker GAD65/67 (**Extended Data Fig. 3a**). Jaccard similarity analysis further confirmed this identity, showing that the terminally differentiated neuronal population more closely resembled cortical cells rather than ventral neuronal types (**Extended Data Fig. 3b**). Both mHD/+ and hHD/+ MAP2-positive neurons showed reduced total dendritic length (TDL) compared to respective wild- type controls (mWT/+ and hWT/+) (**Fig. 3b,c**), with hHD/+ neurons showing the most severe deficit (median TDL: 93.7 μm for hHD/+ versus 130.6 μm for mHD/+; **Fig. 3b,c**). *Htt* deletion (RMCE/-) also caused dendritic shortening, although less severe than hHD/+ (**Fig. 3c**).

**Fig. 3:**
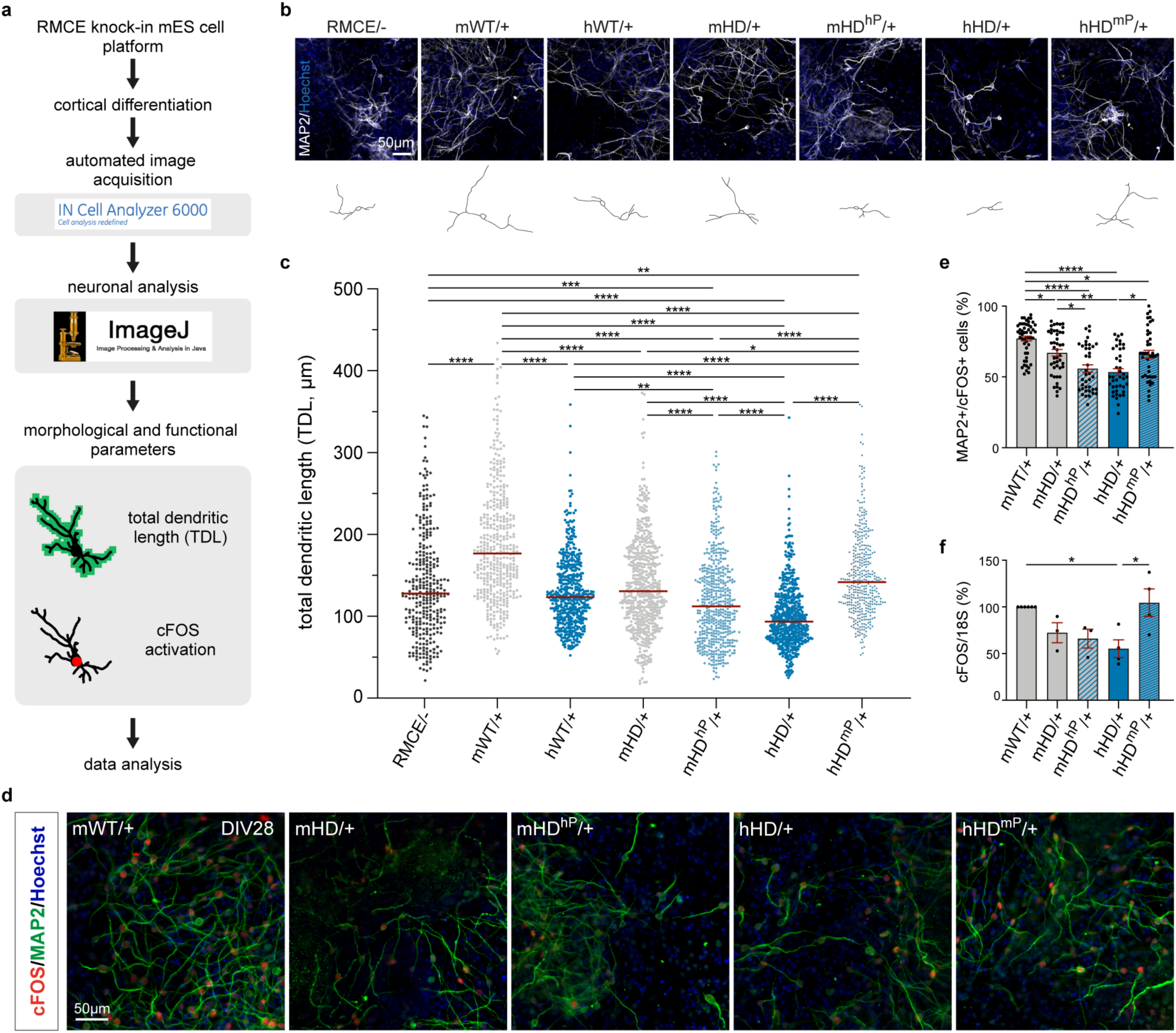
The PRD confers greater pathogenicity to human HD exon 1 in differentiated neurons. **a**, Workflow of morphological and functional analysis on *in vitro* differentiated neurons. **b**, Representative images of neurons derived from mWT/+, mHD/+, mHD^hP^/+, hWT/+, hHD/+, and hHD^mP^/+ mESC lines stained for MAP2 (white) and Hoechst (blue) at DIV28 of differentiation. Scale bar: 50 μm. **c**, Quantification of total dendritic length (TDL) of MAP2+ neurons by ImageJ. Each dot represents the TDL of a single MAP2+ neuron; red line indicates the median of >400 neurons per cell line (n=3 independent replicates). **d**, Representative immunofluorescence images of neurons at DIV28 following a 3-hour glutamate (glu) stimulation, stained for MAP2 (green), cFOS (red), and Hoechst (blue). **e**, Quantification of glu-induced cFOS activation. Each dot represents the % of MAP2+/cFOS+ neurons per field. Data are shown as mean ± SEM from n=3 independent experiments. **f**, qPCR analysis of cFOS expression in glu-stimulated neuronal cultures. Each dot represents an independent replicate, normalized to the mWT/+ condition. Data are shown as mean ± SEM from n≥3 independent experiments. In (**c**) and (**e**) Kruskal-Wallis test with Dunn’s post hoc test: * P<0.05, ** P<0.01, **** P<0.0001. In (**f**) ordinary one-way ANOVA test with Tukey’s post hoc test: * P<0.05.

Functionally, HD neurons are known to exhibit reduced *cFOS* activity^38,39^. Consistent with this, we observed impaired induction of the immediate early gene *cFOS* following glutamate stimulation in our system. Both mHD/+ and hHD/+ neurons displayed a reduced percentage of MAP2+/cFOS+ cells compared to their respective wild-type controls (**Fig. 3d,e**). Notably, hHD/+ neurons showed a ∼20% stronger reduction in *cFOS* activation than mHD/+ neurons (**Fig. 3d,e**). This effect was further confirmed by RT-PCR, which revealed a significant decrease in *cFOS* mRNA levels in hHD/+, while levels remained unchanged in mHD/+ neurons (**Fig. 3f**). Basal *cFOS* expression was comparable across all genotypes **(Extended Data Fig. 4**), indicating that the deficit is stimulus-dependent.

PRD swaps had a clear and reciprocal effect on neuronal phenotypes. Consistent with our observations in neural cysts, mHD^hP^/+ neurons exhibited a more severe reduction in TDL compared to mHD/+ (**Fig. 3b,c**). In contrast, hHD^mP^/+ neurons showed a 51% increase in TDL relative to hHD/+ (**Fig. 3b,c**). Functional measures mirrored these structural changes: cFOS induction following glutamate stimulation was further reduced in mHD^hP^/+ neurons (55.8%, SEM ±2.6) compared to mHD/+ (67.0%, SEM ±2.3), whereas hHD^mP^/+ neurons showed a 23.6% increase in MAP2+/cFOS+ cells relative to hHD/+ (**Fig. 3d,e**). RT-PCR analysis confirmed elevated *cFOS* mRNA levels in hHD^mP^/+ neurons versus hHD/+ cultures (**Fig. 3f**), reinforcing the extent of functional rescue.

Together, these results demonstrate that human HD exon 1 is intrinsically more pathogenic than the murine version, and that this enhanced toxicity is critically encoded within the PRD. These findings establish the PRD as a central determinant of early-to-late HD-related neuronal dysfunction.

### The murine PRD restores the altered proteomic profile of humanized HD neurons

To investigate the molecular pathways involved, we performed global transcriptomic and proteomic profiling of neurons at DIV28 from mWT/+, hHD/+ and hHD^mP^/+ (**Fig. 4a** and **Extended Data Fig. 5a**). Comparative proteomics identified 389 differentially expressed proteins (DEPs) across the three genotypes. Hierarchical clustering grouped hHD^mP^/+ closer to mWT/+ neurons than to hHD/+ neurons, indicating that the murine PRD partially restores the HD proteomic signature (**Fig. 4b**). Cluster analysis revealed six distinct expression patterns (**Fig. 4b**). Clusters 3 and 4 were the most relevant: proteins that were up-regulated (Cluster 3) or down-regulated (Cluster 4) in hHD/+ were restored to near-normal levels in hHD^mP^/+ neurons (**Fig. 4c,d**). Cluster 3 and 4 were also the most abundant – 48 and 263 proteins respectively (i.e., 12.4% and 67.6% of DEPs; **Fig. 4c**) – reaching a total of 311 proteins (80% of DEPs). Clusters 5 and 6 contained proteins whose expression was unchanged or altered independently of PRD status (**Fig. 4c,d**). Due to their small size, clusters 1 and 2 were excluded from further analysis.

**Fig. 4:**
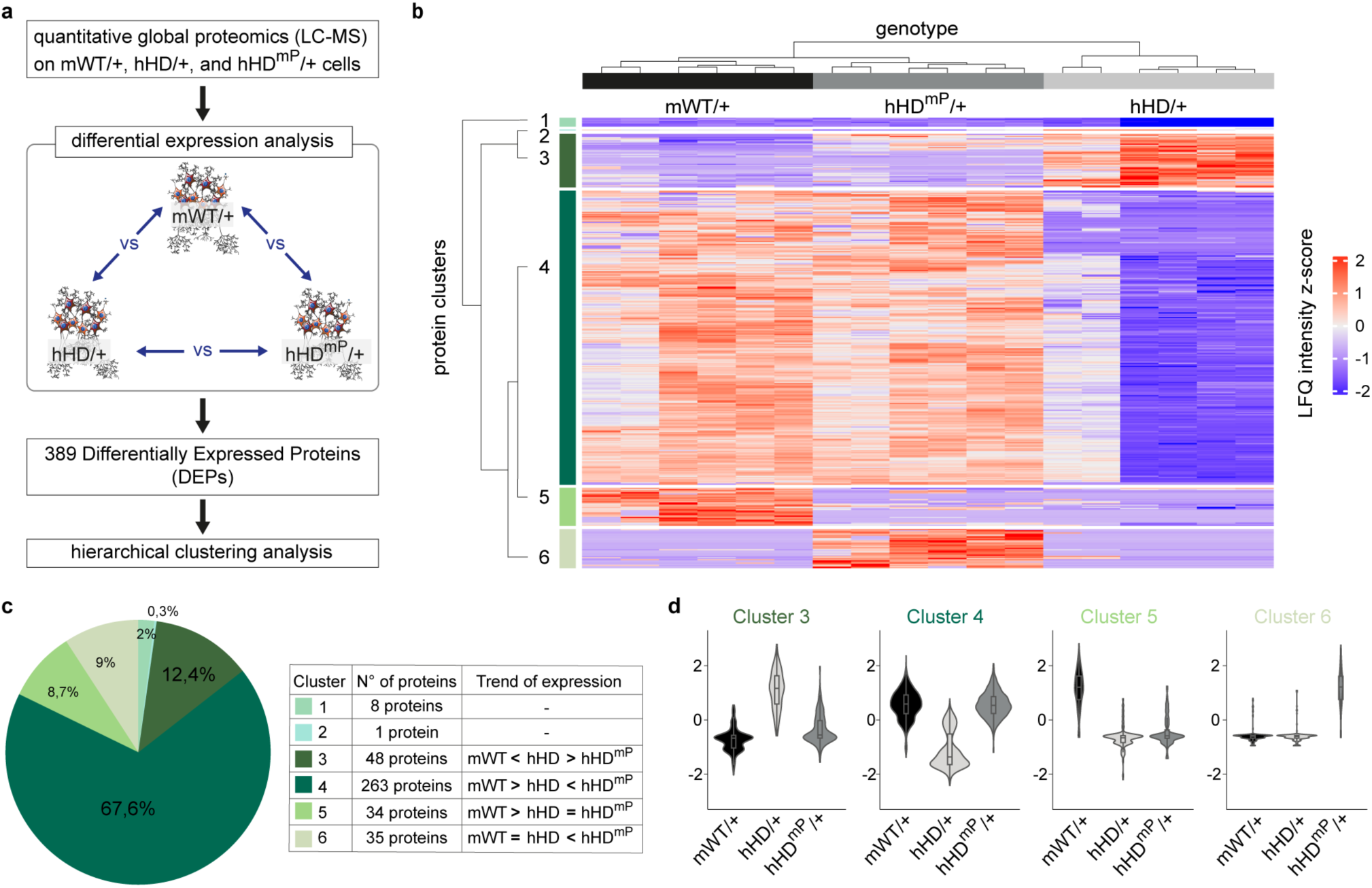
The murine PRD normalizes the proteomic alterations in humanized HD neurons. **a**, Schematic workflow of the proteomic analysis performed on neurons derived from mWT/+, hHD/+, and hHD^mP^/+ mESC lines. **b**, Heatmap showing hierarchical clustering of the 389 DEPs, with expression levels presented as z-scores of label-free quantification (LFQ) intensity values. **c**, Left: pie chart showing the distribution of DEPs across six clusters identified by clustering analysis. Right: table reporting the number of proteins per cluster and their relative expression. **d**, Violin plots illustrating the expression profiles of proteins in the most abundant clusters (Clusters 3 to 6) across the three genotypes.

Transcriptomics identified 2,522 differentially expressed genes (DEGs; **Extended Data Fig. 5a,b**) across genotypes. However, no transcriptional differences were detected between hHD/+ and hHD^mP^/+ neurons (**Extended Data Fig. 5b-d**), indicating that the murine PRD minimally affects gene expression but robustly restores protein-level abnormalities via a post-transcriptional mechanism, likely involving regulation of translation, protein stability, or degradation.

Consistent with the observed transcriptomic changes in hHD/+ neurons, gene ontology (GO) enrichment analysis showed disruption in pathways related to RNA processing, ribosomal function, cilium organization, and intracellular transport (**Extended Data Fig. 5e**). Of the 2,522 DEGs, only 283 overlapped with DEPs, suggesting partial convergence (**Extended Data Fig. 5f**). GO analysis of shared genes/proteins confirmed predominant perturbations in RNA-related pathways (**Extended Data Fig. 5g**).

In summary, replacing the human PRD with the murine sequence is sufficient to restore the pathological proteomic profile of humanized HD neurons to a state resembling that of healthy neurons. However, this substitution does not reverse transcriptional alterations, indicating that the rescue is primarily exerted at the protein level.

### Functional characterization of PRD-rescued proteins reveals the actin cytoskeleton pathway and MKL2/MRTFB as key mediators of phenotypic recovery

To determine which pathways were restored by PRD replacement, we analyzed the 311 rescued proteins from Clusters 3 and 4. GO enrichment revealed roles in transcription, translation, vesicle trafficking, DNA repair, epigenetic, cytoskeleton regulation and morphogenesis (**Extended Data Fig. 6a,b**). STRING network analysis identified three HTT-associated protein clusters (**Extended Data Fig. 7a**). Notably, one of these clusters (Cluster 2) included a sub-network of nine proteins involved in actin cytoskeleton organization. Further supporting this, upstream regulator analysis using Ingenuity Pathway Analysis (IPA) identified 224 predicted regulators of the rescued proteins. All top 10 regulators – ranked by the number of targets – were linked to actin cytoskeleton regulation (**Extended Data Fig. 7b)**, suggesting this pathway plays a dominant role in PRD-mediated rescue. Among these top regulators, HTT itself was identified, further reinforcing the notion that HTT function is intrinsically linked to cytoskeletal regulation. Notably, MKL2 (also known as MRTFB) – an actin-binding transcription coactivator – was among the most significantly rescued proteins, together with SCMH1 and UBE2B (**Fig. 5a**).

**Fig. 5:**
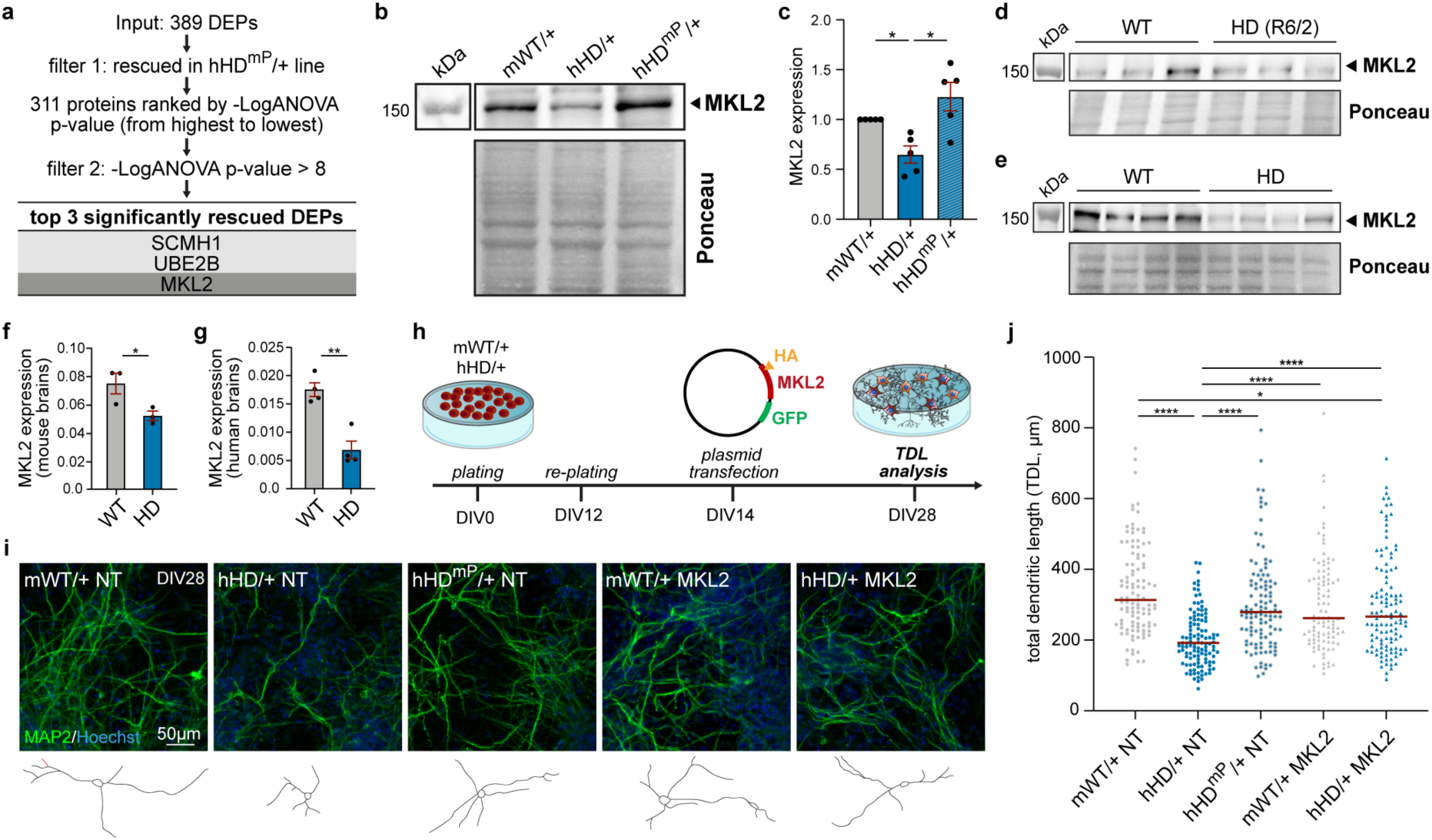
MKL2 is suppressed in the pathological context and its restoration reverses morphological abnormalities in HD neurons. **a**, Schematic of the workflow used to identify the three most significantly rescued proteins. **b**,**c**, Representative western blot image (**b**) and quantification (**c**) of MKL2 expression in neurons derived from mWT/+, hHD/+, and hHD^mP^/+ lines. Total protein stain (ponceau) was used as a loading control. Data are shown as mean ± SEM from n=5 independent experiments. Each dot represents an individual replicate, normalized to the respective mWT/+ condition. One-way ANOVA with Tukey’s post hoc test: * p<0.05. **d**-**g**, Western blot images and quantifications of MKL2 expression in the cortex of the R6/2 HD mouse model (**d**,**f**) and in post-mortem cortical samples from HD patients and controls (**e**,**g**). Total protein stain (ponceau) was used as a loading control. Data are shown as mean ± SEM from n=3 animals (**f**) and n=4 human individuals (**g**) per genotype. Unpaired t-test: * p<0.05, ** p<0.01. **h**, Schematic of MKL2 plasmid transfection during neuronal differentiation. **i**, Representative immunofluorescence images of neurons from mWT/+ NT (Non-Transfected), hHD/+ NT, hHD^mP^/+ NT, mWT/+ MKL2, and hHD/+ MKL2 conditions, stained for MAP2 (green) and Hoechst (blue) at DIV28. Scale bar: 50 μm. **j**, TDL quantification of MAP2+ neurons by ImageJ. Each dot represents the TDL of a single MAP2+ neuron; red line indicates the median. Data from n=4 independent replicates. Kruskal-Wallis test with Dunn’s post hoc test: * P<0.05, **** P<0.0001.

These converging bioinformatic analyses strongly implicate the actin cytoskeleton pathway – and MKL2 in particular – as key contributors to the amelioration of HD-associated defects upon replacement of the human PRD with its murine counterpart.

### MKL2 normalization restores dendritic growth defects in HD neurons

To validate the proteomic findings, we assessed MKL2 expression by western blot in neuronal lysates from five independent differentiations. Consistent with the proteomic data, MKL2 levels were significantly reduced in hHD/+ neurons and restored to normal levels in hHD^mP^/+ neurons (**Fig. 5b,c**). Similar reductions were observed *in vivo*, in the cortex of R6/2 mice compared to wild-type littermates (**Fig. 5d,f**), as well as in post-mortem cortical tissue from HD patients compared to controls (**Fig. 5e,g**). These findings establish MKL2 dysregulation as a consistent and robust feature of HD pathology both *in vitro* and *in vivo*.

To determine whether MKL2 contributes functionally to the rescue of dendritic architecture, we overexpressed HA-tagged MKL2 in maturing neurons (DIV14) using a P2A-GFP construct (**Fig. 5h**). Successful expression was confirmed by detection of both HA-MKL2 and GFP in transfected neurons (**Extended Data Fig. 8a,b**). At DIV28, TDL was measured. As expected, hHD/+ neurons exhibited reduced TDL, which was rescued in hHD^mP^/+ neurons. Importantly, MKL2 overexpression in hHD/+ neurons restored TDL to wild-type levels (**Figure 5i,j**), demonstrating that MKL2 is functionally sufficient to counteract dendritic growth defects in HD neurons.

In conclusion, our findings establish MKL2 plays as a pivotal mediator of the structural abnormalities observed in HD neurons. Its normalization – whether by PRD exchange or direct overexpression – is sufficient to restore dendritic complexity. These results support a model in which the PRD- encoding region of *HTT* contributes to HD pathogenesis, at least in part, by modulating the actin cytoskeleton pathway through regulation of MKL2.

## Discussion

HD is uniquely human, yet it is widely modeled in animals. To date, however, no comparative structure-function studies have systematically explored whether human-specific regions of HTT contributes to pathogenic mechanisms. This represents a significant gap in the HD literature.

In this study, we used mini-organoids (“neural cysts”) and differentiated neurons from our HuntEx1 platform to demonstrate that mutant human *HTT* exon 1 induces more severe phenotypes than its murine equivalent, providing *in vitro* evidence of species-specific pathogenicity. These findings complement *in vivo* studies showing that mice expressing mutant human exon 1 exhibit more pronounced pathology than those expressing the murine version^28^.

Importantly, the more prominent phenotypes observed in hHD/+ versus Htt knock-out lines suggest that the observed defects reflect gain-of-function mechanisms, not just loss of normal huntingtin activity. This interpretation aligns with earlier literature emphasizing the dual contribution of loss- and gain-of-function in HD pathogenesis^2,3,40^.

We further mapped this human-specific toxicity to a discrete element within HTT exon 1 – the PRD – and found that replacing the human with its murine ortholog was sufficient to rescue the pathological phenotype. These data provide direct evidence for the hypothesis that *cis*-elements within the human PRD modulate HD severity, expanding on previous genetic studies suggesting that PRD polymorphisms affect disease outcome^17^. Our neural cyst model also revealed that these PRD- driven effects emerge early in neurogenesis, reinforcing the view that HD has a neurodevelopmental component^29–34^.

The importance of the PRD is supported by evolutionary, biochemical and molecular data. Notably, the PRD co-evolved with the polyQ expansion, suggesting functional interplay^24^. In human populations, CAG and CCG length repeats are genetically correlated^26,27^, suggesting that PRD structure might influence CAG expandability. The PRD also modulates HTT aggregation, solubility and turnover^4,19–21^ and mice with the human exon 1 exhibit increased pathology and toxic *HTT1a* transcript^28^. Indeed, targeting the PRD with intrabodies or deleting it reduces mutant huntingtin toxicity *in vitro* and *in vivo*^22,23^. Taken together, these findings support the hypothesis that CAG tract toxicity and expandability are closely linked to the molecular dynamics and functional properties of the adjacent PRD-encoding region.

We also observed that the detrimental impact of human PRD in neurons manifests predominantly at the protein level, without altering gene transcription, suggesting a PRD-mediated post-transcriptional mechanism involving translational control or altered protein turnover. This was anticipated, given previous reports showing that the HTT PRD is primarily involved in protein binding^41–43^ rather than transcriptional regulation. Indeed, the PRD mediates protein-protein (PP) interactions that underlie numerous cellular processes^44^.

Several molecular mechanisms may underlie the contribution of the PRD to HD pathology. First, given HTT’s established role as a molecular scaffold^2,3^, the PRD may facilitate specific PP interactions. The human PRD may recruit pathogenic interactors that initiate HD-relevant cascades, leading to widespread proteomic alterations. In contrast, the murine PRD may lack these interactions, thereby restoring the proteomic landscape in humanized HD contexts. Notably, the absence of a concomitant transcriptomic rescue suggests that HTT’s transcriptional regulatory function^2,3,45,46^ is not primarily mediated by the PRD. Comparative interactome profiling could help identify these species-specific partners that underlie these differential effects.

A second hypothesis involves previously described phenomena of ribosome stalling and collision at the polyQ–PRD junction of HTT, which lead to ribotoxicity and translational stress^47,48^. Differential levels of stalling across PRDs of distinct origin could thus explain how the mouse PRD enabled the observed rescue at the global proteomic level while leaving transcription unaffected.

Another possibility is that the PRD influences *HTT* splicing, particularly the production of toxic truncated transcripts such as *HTT1a*^49,50^. This is consistent with *in vivo* findings showing increased *HTT1a* levels in mice expressing the human HD exon 1^28^, and our knock-in cell data further support this mechanism (*Maffezzini et al., under revision, co-submission*).

A fourth, non-exclusive hypothesis is that the PRD acts as a *cis*-modifier of somatic CAG instability, a key driver of disease progression^11–13,51^. The co-evolution of the PRD and the polyQ tract, together with the observed CAG-CCG length correlations, strengthens this model. These mechanisms are not-mutually exclusive and may collectively underlie the PRD’s contribution to HD pathogenesis.

Our proteomic analysis revealed that replacing the human PRD with its murine counterpart in hHD/+ neurons restored a wild-type-like proteomic profile, with 311 DEPs that were normalized in this context. Functional enrichment analysis highlighted cytoskeleton organization and dynamics as the most affected. This is in line with recent evidence showing that huntingtin regulates cytoskeletal function in both health and disease^52–54^. Among these rescued proteins, SCMH1, UBE2B, and MKL2 emerged as the most significantly normalized. SCMH1 participates in Polycomb-mediated chromatin remodeling; UBE2B is a key ubiquitin-conjugating enzyme essential for post-replicative DNA repair; and MKL2 is a bifunctional, brain-enriched protein that regulates actin cytoskeleton dynamics and dendritic transcriptional programs, including immediate early genes, like cFOS^55^. Given its central role in cytoskeleton regulation, MKL2 was selected for further investigation. Consistently, MKL2 protein levels were restored in hHD^mP^/+ neurons, and were found to be reduced in both HD mouse models and post-mortem human brain. These data validate MKL2 as a functionally relevant mediator of HD pathology. Moreover, exogenous MKL2 expression in hHD neurons rescued dendritic complexity, supporting a model in which the murine PRD activates a process that restores MKL2 levels, thereby reversing dendritic abnormalities.

Our results are in line with prior studies showing that MKL2 overexpression in primary rat cortical neurons enhances dendritic length and branching^56,57^, whereas MKL2 knock-down or knock-out leads to reduced dendritic outgrowth^56,58–61^.

The precise mechanism by which the murine PRD normalizes MKL2 levels remains unknown. It is plausible that the murine PRD alters direct or indirect interactions between HTT and MKL2 or its adaptors, leading to two non-mutually exclusive outcomes: on one hand, an altered subcellular localization of MKL2 between nucleus and cytoplasm; on the other hand, modulation of MKL2 stability via degradation pathways. Both would result in MKL2 depletion - either at dendrite sites, where it supplies G-actin for F-actin assembly, or in the nucleus, where it co-regulates SRF-target genes (e.g. cFOS, cytoskeleton components). In support, recent evidence shows that huntingtin regulates actin dynamics via the RAC1/LIMK/cofilin axis^62^, which in turn interfaces with the MKL2- SRF^60,63^_._

Overall, our findings demonstrate that the PRD-encoding region of HTT contributes to multiple pathological features associated with the HD mutation – ranging from early morphological abnormalities to late-stage alterations in neuronal properties and the proteomic landscape. Notably, all these phenotypes are fully reversed when the human PRD is replaced by its murine counterpart in the mutant exon 1. In this paradigm, PRD-driven effects are mechanistically linked to MKL2 and other actin cytoskeletal components. We conclude that the PRD modulates HD pathogenicity via a species-specific mechanism involving MKL2-dependent cytoskeleton regulation, identifying a previously unrecognized and potentially actionable therapeutic target in HD.

## Materials and methods

### Mouse Embryonic Stem (mES) cell lines

Genome-edited Recombinase-Mediated Cassette Exchange mES cell lines (RMCE/- and RMCE/+ cells), which were previously generated and characterized^24^ were used as *in vitro* model throughout this study. Both originated from the parental mouse E14 ES cell line^64^. The RMCE/- cell line, which carries the RMCE cassette at the *Htt* exon 1 *locus* on one allele and the deletion of the other *Htt* allele, was used as homozygous Htt knock-out line. The RMCE/+ line, which carries the RMCE cassette at the *Htt* exon 1 *locus* on one allele and a non-edited exon 1 on the other *Htt* allele, was used as master cell line to precisely integrate an array of modified *HTT* exons 1.

### HTT knock-in cell generation (HuntEx1 platform)

A previously generated and characterized RMCE/+ master cell line^24^ was selected and used as original cell line to generate the following knock- in lines: mWT cells (carrying a mouse exon 1 with non-pathogenic 7Q length), mHD cells (carrying a mouse exon 1 with pathogenic 107Q length), mHD^hP^/+ cells (carrying a mouse exon 1 with pathogenic 104Q length but with a human PRD), hWT cells (carrying a human exon 1 with non- pathogenic 20Q length), hHD cells (carrying a human exon 1 with pathogenic 72Q length), hHD^mP^/+ cells (carrying a human exon 1 with pathogenic 72Q length but with a mouse PRD). Taken all together, these cell lines constitute the “HuntEx1” platform. mWT, mHD, mHD^hP^, hWT, hHD, and hHD^mP^ RMCE constructs were here inserted by Flp-mediated recombination. These constructs (synthesized by GenScript and cloned into the pUC57 vector) are, after recombination, able to reconstitute the whole genomic portion that was removed by the PuroRΔTK RMCE cassette. To promote recombination, 2 million RMCE/+ cells were nucleofected using the AmaxaTM mES nucleofector kit (Lonza) with 4 μg RMCE donor construct (either mWT, mHD, mHD^hP^, hWT, hHD, or hHD^mP^) and 4 μg pCAG-Flpe:GFP (Addgene, plasmid #13788). Cells were seeded at low density (4- 8’000 cells/cm^2^) and exposed to 1 μM ganciclovir selection 72h post nucleofection. Recombination efficiency was higher than 60% for all constructs. Clones were individually validated. The first step of clone validation consisted of two PCRs to verify the exchange of the RMCE cassette and the sequence of HTT exon 1 *locus*, using the respective primer pairs. PCR products from the HTT exon 1 *locus* were purified using the QIAquick PCR-purification Kit (QIAGEN) and Sanger sequenced (GATC – Eurofins). The primers used for the PCR of exon 1 are the following: ex1-F (5’- CCCCATTCATTGCCTTGCTG-3’) and ex1-R (5’-GGTCGGTGCAGCGGCTCCT-3’). In the second step, HTT expression was checked by western blot analysis using D7F7 antibody (Cell Signaling Technology), while the expression of pluripotency markers (OCT3/4, SOX2) was confirmed by immunofluorescence analyses in all the cell lines generated.

Each knock-in cell line originates from a pool of at least three validated clones.

### Mouse ES cell culture

Mouse ES cells were maintained in Glasgow minimal essential medium supplemented with 10% heat-inactivated fetal bovine serum (vol/vol, EuroClone, REF ECS0186L), 2 mM l-glutamine, 100 μM non-essential amino acids (Gibco, REF 11140-035), 1 mM sodium pyruvate (Gibco, REF 11360-039), 0.1 mM β-mercaptoethanol (Gibco, REF 31350-010), 100 U/mL penicillin, 100 μg/mL streptomycin (EuroClone, REF ECB3001D) and 1,000 U/mL murine leukemia inhibitor factor (LIF, ESGRO) (Millipore, REF ESG1107), hereafter named “GMEM”. Cells were maintained in 0.2% gelatinized tissue culture flasks and were passaged every 2 days after dissociation with 0.05% trypsin-EDTA (vol/vol) (Gibco, REF 15400-054).

### Neural cyst differentiation

Neural cyst differentiation of our ES cell lines was based on *Meinhardt et al., Stem Cell Reports 2014*^35^. ES cells were dissociated, and 5 x 10^4^ cells were transferred in 30 μL Matrigel (Cultrex BME; Biotechne, REF 3432-010-01). One single Matrigel drop with cells was plated onto each well of an optical 24-multiwell plate. After 15 min at 37°C to allow Matrigel solidification, “modified N2B27” medium was added to the plate. “Modified N2B27” medium is composed of a 1:1 mixture of DMEM/F12 21331 (REF 21331-020) and Neurobasal (REF 21331- 049) medium containing 1:100 B27 supplement (Gibco, REF 17504-044), 0.1 mM β- mercaptoethanol (Gibco, REF 31350-010), and 1:200 modified N2 supplement. This modified N2 was obtained by adding 2 mg/mL human insulin (Sigma, REF I9278) and 7.5 mg/mL bovine serum albumin (BSA; Sigma-Aldrich REF A3059) to standard N2 supplement (Gibco REF 17502-048). Cells were cultured at 37°C for six days, renewing medium every two days. Primitive neural cysts can be appreciated from DIV3. Mature cysts at DIV6 were fixed in 4% PFA for 20 min at room temperature.

### Neuronal cortical differentiation

ES cells were dissociated and plated onto 0.2% gelatin-coated tissue culture dishes at a density of 1 x 10^4^ cells/cm^2^ in GMEM (DIV-1). The day after, GMEM was replaced by “DDM” medium, which consists of DMEM/F12 (REF 21331-020) medium containing 1:100 N2 supplement (Gibco, REF 17502-048), 2 mM l-glutamine, 100 μM non-essential amino acids (Gibco, REF 11140-035), 1 mM sodium pyruvate (Gibco, REF 11360-039), 0.5 mg/mL BSA (Sigma- Aldrich, REF A3059), 0.1 mM β-mercaptoethanol (Gibco, REF 31350-010), 100 U/mL penicillin, 100 μg/mL streptomycin (EuroClone, REF ECB3001D). Cyclopamine (Sigma, REF C4116) was added in DDM from day 2 to day 10 at a final concentration of 1 µM. On day 12-14, cells were dissociated with Accutase (Millipore, REF SF006) and plated on 10 ng/μL Poly-D-Lysin (Sigma Aldrich, REF P6407) and 3 ng/μL Lamin (Thermo Fisher, REF 23017015) coated tissue culture dishes at a density of 8 x 10^4^ cells/cm^2^ in N2B27 medium and allowed to grow for 14-16 days, until DIV28. N2B27 medium consists of a 1:1 mixture of DMEM/F12 and Neurobasal containing 1:200 N2 supplement (Gibco, REF 17502-048), 1:100 B27 supplement (without vitamin A, Gibco, REF 12587-010), 2 mM l-glutamine, 200 μM non-essential amino acids (Gibco, REF 11140-035), 2 mM sodium pyruvate (Gibco, REF 11360-039), 1 mg/mL BSA (Sigma-Aldrich, REF A3059), 0.2 mM β-mercaptoethanol (Gibco, REF 31350-010), 100 U/mL penicillin, 100 μg/mL streptomycin (EuroClone, REF ECB3001D). Medium was renewed every two days. Terminally differentiated cell cultures (DIV28) were fixed in 4% PFA for 20 min at room temperature.

### Genomic DNA extraction

Genomic DNA extraction was performed either using an in house modified Phenol/Chloroform protocol or using a commercial kit (NucleoSpin Tissue® kit - Machery- Nagel, REF 740952). Quality and concentration of all extracted DNA samples were verified on an agarose gel and by spectrometer analysis (NanoDrop 1000 - ThermoScientific).

### RNA extraction (for Real-Time qPCR or RNA-seq analyses)

RNA was isolated using TRIzol reagent according to the manufacturer’s instruction (Life Technologies). RNA was quantified with Nanodrop and then the integrity was evaluated. Potentially contaminating DNA was removed by DNA-free kit (Ambion).

### Real-Time qPCR

RNA was reverse transcribed using the iScript cDNA Synthesis Kit (Bio-Rad). qPCR was performed using the CFX96 Real-Time System (Bio-Rad) and analyzed with the CFX Manager Software (Bio-Rad). All reactions were performed in 15 μL containing 50 ng (for cFOS) and 8.3 ng (for 18S rRNA) cDNA and SsoFast EVAGreen Supermix (Bio-Rad).

### RNA-seq analysis

Bulk RNA sequencing was performed on mWT, hHD and hHD^mP^ cell lines (3 biological replicates per cell line) at the end of neuronal differentiation (DIV28). Stranded polyA libraries were sequenced by the Illumina NovaSeq technology (Eurofins Genomics), with an average depth of 40 million 150 bp paired-end reads per sample.

Sequencing reads were preprocessed with nf-core rnaseq v3.14.0-gb89fac3 Nextflow pipeline^65,66^. Specifically, sequencing reads were trimmed and quality filtered with fastp v0.23.4^67^ and their sequencing quality was checked with FastQC v0.12.1 (https://www.bioinformatics.babraham.ac.uk/projects/fastqc/). Reads generated from residual ribosomal RNA were then identified and removed with SortMeRNA v4.3.4^68^. Filtered reads were aligned to GRCm39 mouse reference genome with STAR v2.7.10a^69^ and the resulting alignments were compressed and saved in sorted BAM files with Samtools v1.17^70^. RSEM v1.3.1^71^ was then used to estimate gene expression levels from BAM files, using annotations from Ensembl v.110 GTF file for GRCm39 reference genome.

Gene counts were imported in R environment with *tximport* function^72^ and a DESeq object was created with *DESeqDataSetFromTximport* function from DESeq2 package v1.12.3^73^. Genes with less than 5 counts in at least three samples were discarded. A differential expression analysis was then performed using *DESeq* function, using GT and Batch as covariates. Genes were considered as differentially expressed (DEGs) in case padj < 0.05 and |log2FC| > 0.25.

The number of DEGs in each comparison (hHD vs mWT, hHD^mP^ vs mWT, hHD vs hHD^mP^) was plotted as a barplot with ggplot2 v3.5.0^74^ and their intersection was plotted using *upset* function from UpSetR v1.4.0 package^75^. Intersection analysis between DEGs and DEPs considered padj < 0.05 for DEG identification and FDR < 0.05 for DEP analysis.

Read counts for DEGs were normalized and transformed with vst function from DESeq2 package and corrected for batch effect using *removeBatchEffect* function from Limma v3.56.2 package^76^. Transformed and batch-corrected counts were scaled by the mean expression value of the gene across samples and saturated between 0.9 and 1.1. Expression values were then plotted as a heatmap with *pheatmap* function from pheatmap v1.0.12 package^77^.

Gene Ontology enrichment analyses were performed for up- and down-regulated DEGs using *enrichGO* function of clusterProfiler v3.0.4 package^78^. Specifically, expressed genes were used as universe, to account for the background, and a p-value threshold of 0.05 was used. Enriched Biological Processes were plotted with *dotplot* function from enrichPlot v1.20.3 package (https://yulab-smu.top/biomedical-knowledge-mining-book/index.html).

#### “Jaccard similarity” index analysis

Read counts from bulk RNA-seq of in vitro samples with mWT/+ genotype were normalized by library size and feature length to obtain TPM and the average expression value for each gene across replicates was calculated. To assess whether the transcriptional profile of mWT/+ in vitro neurons more closely resembled those of in vivo neurons from striatal or cortical origin, read counts from single-nuclear sequencing of three wild-type (mWT/+) mouse fetal samples at embryonic day 15 (E15) from both Lateral Ganglionic Eminence (LGE, the developing striatum) and cortex (CTX) were imported into R environment (unpublished data, *paper under revision*). For each gene and sample, read counts were collapsed across cells to obtain a pseudobulk transcriptional profile for each sample with collapse counts function from presto v1.0.0 package (immunogenomics/presto: Fast Wilcoxon and auROC; https://github.com/immunogenomics/presto). Counts were then normalized by library size and feature length to obtain TPM and the average expression value for each gene and tissue was calculated. Tissue-specific genes, showing no read counts at all in any of the three replicates in one but not the other tissue were identified. The expression profile of tissue-specific genes was compared pairwise across conditions (in vitro, in vivo LGE, in vivo CTX) by using the Jaccard Similarity Index and was represented as a heatmap using pheatmap v1.0.12 package^77^.

### Protein lysates preparation and western blot

Neuronal pellets and cortex samples were lysed in RIPA buffer (50 mM Tris-HCl pH 8, 150 mM NaCl, 0.1% SDS, 1% nonidet P40, 0.5% sodium deoxycholate, wt/vol) with 1 mM PMSF and protease inhibitor (Thermo Scientific, REF 1861281). Lysates were cleared by centrifugation at 12,000g and 4 °C for 30 min. The resulting supernatant was collected. Protein concentration was determined with the Pierce-BCA Protein assay kit (Thermo Scientific, REF 23225). Protein extracts (30 μg) were loaded on SDS-PAGE gels, with a polyAcrylamide concentration depending on the protein of interest. Separated proteins were transferred to a nitrocellulose membrane and stained with total protein stain (Ponceau S, Sigma-Aldrich P7170). The membranes were blocked after washing with 5% BSA (wt/vol, Biorad, REF 170- 6404) in Tris-buffered saline (TBS) and 0.1% Tween-20 (vol/vol), and incubated with primary antibody at room temperature for 3 h. After washing, filters were incubated for 1 h at room temperature with a secondary antibody (peroxidase conjugate, Biorad, 1:3000) and then washed three times with TBS and 0.1% Tween-20. Clarity Western ECL Substrate (Biorad, REF 170-5061) was used to visualize immunoreactive bands by chemiluminescence detection on a ChemiDoc MP Imaging System (Biorad).

Mouse cortex (8 weeks) samples from wild-type and R6/2 animals (B6CBAF1/J background) were purchased from Jackson Laboratory. All protocols involving animals were carried out in accordance with institutional guidelines in compliance with Italian law (D. Lgs no. 2014/26, implementation of the 2010/63/UE; authorizations 686/2021-PR, issued on September 6th, 2021). Human post-mortem cortex samples (Brodmann’s area 9) were obtained from the Harvard Brain Tissue Resource Center (HBTRC) (Belmont, Massachusetts, USA) under Ethics Committee 18.12.13, Ethical Approval 74/14.

### Global proteomics

#### Cell lysis

Neuronal pellets (DIV28) were resuspended in 8 M urea and 20 mM Tris*HCl pH 8 (Urea buffer) and sonicated in the Bioruptor at 4 °C for 3 cycles, alternating between 30 s of sonication and 30 s of idle time. Cell lysates were centrifuged for 15 min at 14000 rcf and the supernatants were collected and stored at −80°C until required. An aliquot was used to estimate protein concentration by BCA assay (Thermo Scientific).

*Proteins digestion and peptides purification.* 50 μg of proteins were in-solution digested according to the following protocol: cysteines reduction and alkylation were performed adding both tris (2- carboxyethyl) phosphine (TCEP - Thermo Scientific) at 10 mM final concentration and 2- Chloroacetamide (CAM - Sigma-Aldrich) at 40 mM final concentration. Samples were incubated for 30 min at room temperature. Then, samples were diluted with 100 mM Tris*HCl pH 8 to reach 5M urea and proteins were digested first with Lys-C (Lysyl Endopeptidase, Mass Spectrometry Grade – Wako) for 1h at RT in a 1:30 enzyme:protein ratio; then, after a further dilution with 100 mM Tris*HCl pH 8 to 2M urea, proteins were digested with trypsin (Sequencing Grade, modified – Roche) overnight at room temperature in a 1:30 enzyme:protein ratio. Samples were acidified to 1% with formic acid and desalted and concentrated passing through the C18 StageTip plugs^79^. The eluted peptides were dried and resuspended in 15 μl of 5% formic acid.

#### Nanoflow LC MS/MS Analysis

1.5 μl of each digested sample were analyzed as technical replicate on a UPLC easy-nLC 1200 coupled with a Quadrupole Orbitrap Exploris 480 mass spectrometer equipped with a Nano Flex ESI-source and the FAIMS Pro interface operating at CV -50/-70 (Thermo Fisher Scientific). The nLC system was connected to a 25 cm fused-silica emitter of 75 μm inner diameter (New Objective), packed in house with ReproSil-Pur C18-AQ 1.9 μm beads (Dr. Maisch) using a high-pressure bomb loader (Proxeon). Peptide separation was achieved on a linear gradient of 131 min, starting from 97% of solvent A (0.1% formic acid, 2% acetonitrile) for 1min and ramping to 19% of solvent B (0.1% formic acid, 80% acetonitrile) in 72 min, then to 29% of B in 28 min and to 41 % of B in 20 min. This was followed by a washout step of 3 min increase to 95% of B which was held for 7 min. Flow rate was kept at 0.200 μl min-1. Spray voltage was set to 2.2 kV, funnel RF level at 40, and heated capillary at 300°C. MS data were acquired in positive data-dependent acquisition mode (DDA), with a top 25 method for HCD fragmentation. Full MS spectra (300–15000 Th) were acquired in the Orbitrap with 60,000 resolution, Time 100 ms, Normalized AGC target 100%, Microscan 1, AGC target Custom, Maximum IT Custom, RF Lens 40%. For fragment spectra the resolution was set to 15,000, Normalized AGC target 100%, Microscan 1, AGC target Custom, Maximum IT Auto; normalized collision energy 28%, isolation width 1.6 m/z with an exclusion duration of 20 s.

#### Data processing and label-free quantification

Raw data were processed using MaxQuant (ver. 2.0.3.0)^80^ with default setting, searching against the Database uniprot_cp_Mouse_2020 containing also the sequences of Htt hHDQ/+ (G11111) and Htt hHDmP/+ (G11112), in which trypsin enzyme was selected with up two missed cleavages. Cysteine Carbamidomethylation was used as a fixed modification, Methionine Oxidation and Protein N-terminal Acetylation as variable modifications. Mass deviation for MS and MS/MS peaks were set at 10 ppm and 20 ppm respectively. The peptide and protein false-discovery-rates (FDRs) were set to 0.01 with a minimal peptide length of seven amino acids; for high-confidence protein identification a minimum of two peptides and at least one unique peptide were required.

Label-free analysis was carried out, including the ‘match between runs’ option (time window of 2 min). The lists of identified proteins were filtered to eliminate reverse hits and known contaminants. Statistical analyses were done using the Perseus program (ver. 1.6.2.3)^81^ in the MaxQuant environment, considering the protein LFQ intensity normalized based on the z-score; the function ‘imputation’ was selected to replace missing values by random numbers drawn from a normal distribution. Hierarchical Clustering analysis was done applying Anova, Benjamini Hochberg test and an FDR of 0.01.

Proteomics data associated with this paper have been uploaded to the PRIDE repository (https://www.ebi.ac.uk/pride/) under the project accession identifier PXD061056.

#### Bioinformatic analyses

Heatmap representing hierarchical clustering analysis based on LFQ intensity z-score values was generated by pheatmap R package. Then, the analyses focused on differentially expressed proteins (DEPs) that suggested a rescued phenotype, i.e. those showing a similar expression level in wild-type and in human HD exon 1 with a mouse PRD context, as opposed to the fully human HD exon 1 context. To further characterize these 311 “rescued” DEPs, the TopGO R package was used to perform gene ontology enrichment analysis, highlighting the main biological processes in which they are involved. Next, we used the STRING database to identify and visualize networks of protein-protein interactions between members of each cluster, focusing only on validated interactions from experiments and databases with an interaction score greater than 0.45. While it was not among the rescued DEPs, Htt was included in the analysis to identify potentially meaningful direct interactors. Finally, we used the QIAGEN Ingenuity Pathway Analysis (IPA) database to identify the upstream regulators that might explain the observed protein expression changes, as well as to further validate the enrichment and network analyses.

### Electron microscopy

#### TEM sample preparation and imaging

Neural cysts were fixed in 2.5% glutaraldehyde and 2% paraformaldehyde (Electron Microscopy Sciences) in 0.15 M sodium cacodylate buffer (pH 7.4). Samples were washed with 0.1 M cold sodium cacodylate buffer and post-fixed for 1 h on ice in reduced osmium (2% OsO₄ and 1.5% potassium ferrocyanide in 0.15 M cacodylate buffer). After washing with ddH₂O, samples were incubated in 1% thiocarbohydrazide (TCH) for 20 min, rinsed, and treated with 2% OsO₄ in ddH₂O for 30 min at room temperature. Following multiple ddH₂O washes, samples were incubated in 1% aqueous uranyl acetate overnight at 4 °C, then stained with Walton’s lead aspartate for 30 min at 60 °C. Dehydration was performed through an ethanol series, followed by a 10 min incubation in ice-cold anhydrous acetone. Infiltration was carried out using acetone/Epon812 resin (3:1 for 2 h, then 1:1 overnight), followed by 2 h in pure resin and polymerization at 60 °C for 48 h. Ultrathin sections were cut using an ultramicrotome (UC7, Leica Microsystems, Vienna, Austria), placed on transmission electron microscopy (TEM) copper grids and imaged by a Tecnai G2 Spirit microscope (FEI, Eindhoven, The Netherlands) operated at an acceleration voltage of 120 kV, equipped with a S-twin objective lens and a bottom-mount 11MP Orius SC1000 CCD camera (Gatan, Pleasanton, USA).

#### SEM sample preparation and imaging

Neural cysts were fixed in 2% glutaraldehyde in 0.1 M sodium cacodylate buffer for 24 h at 4 °C, then washed three times in the same buffer. Post-fixation was performed in 2% osmium tetroxide in 0.1 M sodium cacodylate for 1 h at room temperature, followed by two rinses in bi-distilled water. Samples were dehydrated through a graded ethanol series (20% to 100%), then treated with 50% and 100% hexamethyldisilazane (HMDS), and air-dried overnight. Neural cysts were opened using 0.5 mm tungsten needles under a Leica stereomicroscope and mounted on 12 mm-sized aluminium stubs. Prior to imaging, samples were coated with a 20 nm platinum-palladium layer using a 208HR (Cressington, Watford, UK) sputter coater with an Ar- plasma beam current of 40 mA. Scanning electron microscopy (SEM) imaging was performed by a Merlin Field Emission (FE)-SEM (Zeiss, Oberkochen, Germany) operating at an acceleration voltage of 5 kV with a 200 pA electron beam current and 10 mm of working distance, collecting the secondary electrons signal by an in-chamber Everhart-Thornley detector.

#### Immunofluorescence

Neural cysts were blocked for 1 hr at room temperature (1 × PBS, 0.5% BSA, 0.3% Triton X-100) after quenching (1 × PBS, 200 mM glycine, 0.3% Triton X-100). ES cells and neurons, were permeabilized and blocked in blocking buffer containing PBS with 0.5% Triton X- 100 (vol/vol) and with 5% fetal bovine serum (vol/vol) for 1 h. Primary antibodies were diluted in the respective blocking buffer and applied overnight at 4 °C. After three washes in PBS, appropriate secondary antibodies conjugated to Alexa fluorophores 488 or 568 or 647 (Molecular Probes, Invitrogen) and diluted 1:500 in blocking solution, were applied for 1 h at room temperature. Cells were incubated for 10 min with Hoechst 33258 (5 μg/mL, Molecular Probes, Invitrogen). Images were acquired with an IN Cell Analyzer 6000 (GE Healthcare Life Sciences) and processed with the software ImageJ (US National Institutes of Health) and CellProfiler (vers.2.1.1). For neural cysts, 10 z-stacks per field (step-size 10μm) were acquired with “Max intensity projection” mode.

### Multiparametric quantification of neural cysts and neurons

#### Morphological neural cyst quantification

Three wells per cell line were analyzed per experiment, with 8 randomly selected fields per well, leading to a total of around 2,000–3,000 neural cysts. They were blind quantified on DIV6 of neural differentiation after staining for NESTIN and PALS1. CellProfiler software was used to automatically quantify “neural cyst” objects measuring 46 size and shape parameters for each neural cyst (object). Details on the CellProfiler quantification pipeline are reported in **Extended Data Fig. 2d**. Analysis from data tables were handled with custom Matlab® pipelines. Cells with unusual shapes were identified from the area vs perimeter plot. This was indeed showing isolated data points that were corresponding to cysts cut by the image borders. These “cyst” objects were automatically identified and discarded from the analysis. We also used a threshold (1000 px^2^) on the area to remove small dimension debris from the dataset. All the parameters were normalized to the 0-1 range and principal component analysis was performed. The PCA analyses naturally separated size and shape parameters pretty well, identifying them as the most important contributors to the first (PC1) and second (PC2) components, respectively.

#### Morphological neuron quantification

For each cell line, the measurements of the total dendritic length were performed on 30 randomly selected fields per experiment. In total, approximately 400 MAP2+ neurons were analyzed per cell line. They were blind quantified on DIV28 of neuronal differentiation after staining for MAP2. Total dendritic length for each neuron was manually measured using the ’Freehand Line’ tool in ImageJ software.

#### cFOS neuron quantification

For each cell line, 20 randomly selected fields were analyzed per experiment. Neurons upon glutamate stimulus (3 hours, 25 mM) or in basal condition (no glutamate) were blind quantified after neuronal differentiation (DIV28) on MAP2 and cFOS staining. CellProfiler software was used to automatically determine the number of cFOS-positive cells, expressed as a percentage of the total Hoechst-positive cell count.

#### MKL2 plasmid transfection

MKL2 plasmid was generated by Genscript, inserting HA-tagged MKL2 isoform 1 (CDS from NCBI Reference Sequence, NM_153588.3) into pcDNA3.1(+)-P2A- eGFP vector.

DNA transfection with the MKL2 plasmid (4 μg) was performed two days after re-plating (DIV12), during the neuronal differentiation protocol using Lipofectamine 3000 (Thermo Fisher Scientific, REF L3000015). Western blot was conducted to assess exogenous MKL2 expression two days after transfection.

#### Statistics for neural cyst and neuron assays

Data from the quantification of our neural cyst assay was analyzed with Kruskal Wallis test and Bonferroni correction. Data from the quantification of neurons was analyzed with ordinary one-way ANOVA or Kruskal Wallis test by using GraphPad Prism (version 10.0.2). Multiple comparisons between cell lines were performed by applying Tukey’s or Dunn’s post-hoc tests.

## Data availability

Fastq files for RNA-sequencing data have been submitted to NCBI SRA (BioProject PRJNA1232624). Proteomics data associated with this paper have been uploaded to the PRIDE repository (https://www.ebi.ac.uk/pride/) under the project accession identifier PXD061056.

## Acknowledgements

We thank Graziano Martello (Department of Molecular Medicine, University of Padova) for E14 cells and advice for their growth; Chiara Cordiglieri (INGM Imaging facility) for her support with imaging acquisition. This project was funded by CHDI Foundation, Inc. New York, (JSC A11103) a nonprofit biomedical research organization exclusively dedicated to developing therapeutics that will substantially improve the lives of HD-affected individuals. In particular, we would like to acknowledge the involved CHDI members for their great support and fruitful scientific discussions.

## Author contributions

RI and EC conceived the study, with the contribution of CM, CT and CZ; RI, CM, AS performed the molecular and cellular biology experiments; RI developed the neural cysts assay and the related multi-parametric quantification pipeline; ECam performed the statistical analysis of the neural cysts dataset; EV and AF performed the electron microscopy imaging; RI and AS performed neuronal phenotypical characterization; RI collected samples for -omics analyses; RI, CM, SM, EC, performed all RNAseq work, including strategy design, RNAseq data analysis and interpretation; RI, CM, AC, AM, AB, EC, performed all proteomics work, including strategy design, proteomics data analysis and interpretation; RI designed the experimental strategy to test MKL2 role; CM and CZ contributed to MKL2 validation and functional experiments. RI assembled the figures and wrote the paper with CM and EC. The manuscript was then read, commented and reviewed by all authors. DPF and TFV provided scientific input. EC coordinated the study, secured the funding and ensured that the descriptions contained in the final version of the paper are accurate and agreed by all authors. The contributions in each section of this paragraph reflect the order of the authors.

## Ethics declarations

Mouse samples were purchased from Jackson Laboratory and all protocols were carried out in accordance with institutional guidelines, in compliance with Italian law (D. Lgs no. 2014/26, implementation of the 2010/63/UE; authorizations 686/2021-PR, issued on September 6th, 2021). Human post-mortem samples were obtained from the Harvard Brain Tissue Resource Center (HBTRC, Belmont, Massachusetts, USA) under Ethics Committee 18.12.13, Ethical Approval 74/14.

## References

1. Macdonald, M. A novel gene containing a trinucleotide repeat that is expanded and unstable on Huntington’s disease chromosomes. Cell 72, 971–983 (1993).

2. Zuccato, C., Valenza, M. & Cattaneo, E. Molecular Mechanisms and Potential Therapeutical Targets in Huntington’s Disease. Physiol. Rev. 90, 905–981 (2010).

3. Saudou, F. & Humbert, S. The Biology of Huntingtin. Neuron 89, 910–926 (2016).

4. Pigazzini, M. L., Lawrenz, M., Margineanu, A., Kaminski Schierle, G. S. & Kirstein, J. An Expanded Polyproline Domain Maintains Mutant Huntingtin Soluble in vivo and During Aging. Front. Mol. Neurosci. 14, (2021).

5. Lee, J.-M. et al. CAG repeat expansion in Huntington disease determines age at onset in a fully dominant fashion. Neurology 78, 690–695 (2012).

6. Keum, J. W. et al. The HTT CAG-Expansion Mutation Determines Age at Death but Not Disease Duration in Huntington Disease. Am. J. Hum. Genet. 98, 287–298 (2016).

7. Djoussé, L. et al. Interaction of normal and expanded CAG repeat sizes influences age at onset of Huntington disease. Am. J. Med. Genet. A. **119A**, 279–282 (2003).

8. Wexler, N. S. Venezuelan kindreds reveal that genetic and environmental factors modulate Huntington’s disease age of onset. Proc. Natl. Acad. Sci. 101, 3498–3503 (2004).

9. Gusella, J. F. & MacDonald, M. E. Huntington’s disease: the case for genetic modifiers. Genome Med. 1, 80 (2009).

10. Pengo, M. & Squitieri, F. Beyond CAG Repeats: The Multifaceted Role of Genetics in Huntington Disease. Genes 15, 807 (2024).

11. Ciosi, M. et al. A genetic association study of glutamine-encoding DNA sequence structures, somatic CAG expansion, and DNA repair gene variants, with Huntington disease clinical outcomes. EBioMedicine 48, 568–580 (2019).

12. Lee, J.-M. et al. CAG Repeat Not Polyglutamine Length Determines Timing of Huntington’s Disease Onset. Cell 178, 887–900.e14 (2019).

13. Wright, G. E. B. et al. Length of Uninterrupted CAG, Independent of Polyglutamine Size, Results in Increased Somatic Instability, Hastening Onset of Huntington Disease. Am. J. Hum. Genet. 104, 1116–1126 (2019).

14. Lee, J.-M. et al. Genetic modifiers of Huntington disease differentially influence motor and cognitive domains. Am. J. Hum. Genet. 109, 885–899 (2022).

15. Lee, J.-M., MacDonald, M. E. & Gusella, J. F. Inherited HTT CAG repeat length does not have a major impact on Huntington disease duration. Am. J. Hum. Genet. 109, 1338–1340 (2022).

16. Hong, E. P. et al. Huntington’s Disease Pathogenesis: Two Sequential Components. J. Huntingt. Dis. 10, 35–51 (2021).

17. Dawson, J. et al. A probable cis-acting genetic modifier of Huntington disease frequent in individuals with African ancestry. Hum. Genet. Genomics Adv. 3, 100130 (2022).

18. Lee, J.-M. et al. Genetic modifiers of somatic expansion and clinical phenotypes in Huntington’s disease highlight shared and tissue-specific effects. Nat. Genet. 57, 1426–1436 (2025).

19. Duennwald, M. L., Jagadish, S., Giorgini, F., Muchowski, P. J. & Lindquist, S. A network of protein interactions determines polyglutamine toxicity. Proc. Natl. Acad. Sci. 103, 11051–11056 (2006).

20. Dehay, B. & Bertolotti, A. Critical Role of the Proline-rich Region in Huntingtin for Aggregation and Cytotoxicity in Yeast. J. Biol. Chem. 281, 35608–35615 (2006).

21. Falk, A. S. et al. Structural Model of the Proline-Rich Domain of Huntingtin Exon-1 Fibrils. Biophys. J. 119, 2019–2028 (2020).

22. Southwell, A. L. et al. Intrabodies Binding the Proline-Rich Domains of Mutant Huntingtin Increase Its Turnover and Reduce Neurotoxicity. J. Neurosci. 28, 9013–9020 (2008).

23. Brady, S. T. et al. Toxic effects of mutant huntingtin in axons are mediated by its proline-rich domain. Brain 147, 2098–2113 (2024).

24. Iennaco, R. et al. The evolutionary history of the polyQ tract in huntingtin sheds light on its functional pro-neural activities. Cell Death Differ. 29, 293–305 (2022).

25. Neema, M. et al. Mutant Huntingtin Drives Development of an Advantageous Brain Early in Life: Evidence in Support of Antagonistic Pleiotropy. Ann. Neurol. 96, 1006–1019 (2024).

26. Squitieri, F. et al. DNA haplotype analysis of Huntington disease reveals clues to the origins and mechanisms of CAG expansion and reasons for geographic variations of prevalence. Hum. Mol. Genet. 3, 2103–2114 (1994).

27. Morovvati, S., Nakagawa, M., Osame, M. & Karami, A. Analysis of CCG Repeats in Huntingtin Gene among HD Patients and Normal Populations in Japan. Arch. Med. Res. 39, 131–133 (2008).

28. Franich, N. R. et al. Phenotype onset in Huntington’s disease knock-in mice is correlated with the incomplete splicing of the mutant huntingtin gene. J. Neurosci. Res. 97, 1590–1605 (2019).

29. Conforti, P. et al. Faulty neuronal determination and cell polarization are reverted by modulating HD early phenotypes. Proc. Natl. Acad. Sci. 115, (2018).

30. Wiatr, K., Szlachcic, W. J., Trzeciak, M., Figlerowicz, M. & Figiel, M. Huntington Disease as a Neurodevelopmental Disorder and Early Signs of the Disease in Stem Cells. Mol. Neurobiol. 55, 3351–3371 (2018).

31. Haremaki, T. et al. Self-organizing neuruloids model developmental aspects of Huntington’s disease in the ectodermal compartment. Nat. Biotechnol. 37, 1198–1208 (2019).

32. Barnat, M. et al. Huntington’s disease alters human neurodevelopment. Science 369, 787–793 (2020).

33. Van Der Plas, E., Schultz, J. L. & Nopoulos, P. C. The Neurodevelopmental Hypothesis of Huntington’s Disease. J. Huntingt. Dis. 9, 217–229 (2020).

34. Galimberti, M. et al. Huntington’s disease cellular phenotypes are rescued non-cell autonomously by healthy cells in mosaic telencephalic organoids. Nat. Commun. 15, (2024).

35. Meinhardt, A. et al. 3D Reconstitution of the Patterned Neural Tube from Embryonic Stem Cells. Stem Cell Rep. 3, 987–999 (2014).

36. Lo Sardo, V., et al. An evolutionary recent neuroepithelial cell adhesion function of huntingtin implicates ADAM10-Ncadherin. Nat. Neurosci. 15, 713–721 (2012).

37. Gaspard, N. et al. An intrinsic mechanism of corticogenesis from embryonic stem cells. Nature 455, 351–357 (2008).

38. Zhang, J. et al. c-fos regulates neuronal excitability and survival. Nat. Genet. 30, 416–420 (2002).

39. Wacker, J. L. et al. Loss of Hsp70 Exacerbates Pathogenesis But Not Levels of Fibrillar Aggregates in a Mouse Model of Huntington’s Disease. J. Neurosci. 29, 9104–9114 (2009).

40. Jiang, A., Handley, R. R., Lehnert, K. & Snell, R. G. From Pathogenesis to Therapeutics: A Review of 150 Years of Huntington’s Disease Research. Int. J. Mol. Sci. 24, 13021 (2023).

41. Harjes, P. & Wanker, E. E. The hunt for huntingtin function: interaction partners tell many different stories. Trends Biochem. Sci. 28, 425–433 (2003).

42. Qin, Z.-H. et al. Huntingtin Bodies Sequester Vesicle-Associated Proteins by a Polyproline- Dependent Interaction. J. Neurosci. 24, 269–281 (2004).

43. Gao, Y.-G. et al. Structural Insights into the Specific Binding of Huntingtin Proline-Rich Region with the SH3 and WW Domains. Structure 14, 1755–1765 (2006).

44. Umumararungu, T. et al. Proline, a unique amino acid whose polymer, polyproline II helix, and its analogues are involved in many biological processes: a review. Amino Acids 56, (2024).

45. Malla, B., Guo, X., Senger, G., Chasapopoulou, Z. & Yildirim, F. A Systematic Review of Transcriptional Dysregulation in Huntington’s Disease Studied by RNA Sequencing. Front. Genet. 12, (2021).

46. Gu, X. et al. Uninterrupted CAG repeat drives striatum-selective transcriptionopathy and nuclear pathogenesis in human Huntingtin BAC mice. Neuron 110, 1173–1192.e7 (2022).

47. Huter, P. et al. Structural Basis for Polyproline-Mediated Ribosome Stalling and Rescue by the Translation Elongation Factor EF-P. Mol. Cell 68, 515–527.e6 (2017).

48. Aviner, R. et al. Polyglutamine-mediated ribotoxicity disrupts proteostasis and stress responses in Huntington’s disease. Nat. Cell Biol. 26, 892–902 (2024).

49. Sathasivam, K. et al. Aberrant splicing of *HTT* generates the pathogenic exon 1 protein in Huntington disease. Proc. Natl. Acad. Sci. 110, 2366–2370 (2013).

50. Neueder, A. et al. The pathogenic exon 1 HTT protein is produced by incomplete splicing in Huntington’s disease patients. Sci. Rep. 7, (2017).

51. Scahill, R. I. et al. Somatic CAG repeat expansion in blood associates with biomarkers of neurodegeneration in Huntington’s disease decades before clinical motor diagnosis. Nat. Med. 31, 807–818 (2025).

52. Tousley, A. et al. Huntingtin associates with the actin cytoskeleton and α-actinin isoforms to influence stimulus dependent morphology changes. PLOS ONE 14, e0212337 (2019).

53. Taran, A. S., Shuvalova, L. D., Lagarkova, M. A. & Alieva, I. B. Huntington’s Disease—An Outlook on the Interplay of the HTT Protein, Microtubules and Actin Cytoskeletal Components. Cells 9, 1514 (2020).

54. Wu, J. The Roles of Cytoskeleton in Huntington’s Disease. in Proceedings of the 1st International Conference on Health Big Data and Intelligent Healthcare 572–576 (SCITEPRESS - Science and Technology Publications, Sanya, China, 2022). doi:10.5220/0011374400003438.

55. Tabuchi, A. & Ihara, D. Regulation of Dendritic Synaptic Morphology and Transcription by the SRF Cofactor MKL/MRTF. Front. Mol. Neurosci. 14, (2021).

56. Ishikawa, M. et al. Involvement of the Serum Response Factor Coactivator Megakaryoblastic Leukemia (MKL) in the Activin-regulated Dendritic Complexity of Rat Cortical Neurons*. J. Biol. Chem. 285, 32734–32743 (2010).

57. Ishibashi, Y. et al. Expression of SOLOIST/MRTFB i4, a novel neuronal isoform of the mouse serum response factor coactivator myocardin-related transcription factor-B, negatively regulates dendritic complexity in cortical neurons. J. Neurochem. 159, 762–777 (2021).

58. O’Sullivan, N. C., Pickering, M., Di Giacomo, D., Loscher, J. S. & Murphy, K. J. Mkl Transcription Cofactors Regulate Structural Plasticity in Hippocampal Neurons. Cereb. Cortex 20, 1915–1925 (2010).

59. Lee, S.-M., Vasishtha, M. & Prywes, R. Activation and Repression of Cellular Immediate Early Genes by Serum Response Factor Cofactors. J. Biol. Chem. 285, 22036–22049 (2010).

60. Mokalled, M. H., Johnson, A., Kim, Y., Oh, J. & Olson, E. N. Myocardin-related transcription factors regulate the Cdk5/Pctaire1 kinase cascade to control neurite outgrowth, neuronal migration and brain development. Development 137, 2365–2374 (2010).

61. Kaneda, M. et al. Synaptic localisation of SRF coactivators, MKL1 and MKL2, and their role in dendritic spine morphology. Sci. Rep. 8, (2018).

62. Wennagel, D., Braz, B. Y., Capizzi, M., Barnat, M. & Humbert, S. Huntingtin coordinates dendritic spine morphology and function through cofilin-mediated control of the actin cytoskeleton. Cell Rep. 40, 111261 (2022).

63. Olson, E. N. & Nordheim, A. Linking actin dynamics and gene transcription to drive cellular motile functions. Nat. Rev. Mol. Cell Biol. 11, 353–365 (2010).

64. Hooper, M., Hardy, K., Handyside, A., Hunter, S. & Monk, M. HPRT-deficient (Lesch–Nyhan) mouse embryos derived from germline colonization by cultured cells. Nature 326, 292–295 (1987).

65. Di Tommaso, P. et al. Nextflow enables reproducible computational workflows. Nat. Biotechnol. 35, 316–319 (2017).

66. Harshil Patel et al. nf-core/rnaseq: nf-core/rnaseq v3.17.0 - Neon Newt. Zenodo 10.5281/ZENODO.13986791 (2024).

67. Chen, S., Zhou, Y., Chen, Y. & Gu, J. fastp: an ultra-fast all-in-one FASTQ preprocessor. Bioinformatics 34, i884–i890 (2018).

68. Kopylova, E., Noé, L. & Touzet, H. SortMeRNA: fast and accurate filtering of ribosomal RNAs in metatranscriptomic data. Bioinformatics 28, 3211–3217 (2012).

69. Stark, R., Grzelak, M. & Hadfield, J. RNA sequencing: the teenage years. Nat. Rev. Genet. 20, 631–656 (2019).

70. Li, H. et al. The Sequence Alignment/Map format and SAMtools. Bioinformatics 25, 2078– 2079 (2009).

71. Li, B. & Dewey, C. N. RSEM: accurate transcript quantification from RNA-Seq data with or without a reference genome. BMC Bioinformatics 12, (2011).

72. Soneson, C., Love, M. I. & Robinson, M. D. Differential analyses for RNA-seq: transcript-level estimates improve gene-level inferences. F1000Research 4, 1521 (2016).

73. Love, M. I., Huber, W. & Anders, S. Moderated estimation of fold change and dispersion for RNA-seq data with DESeq2. Genome Biol. 15, (2014).

74. Wickham, H. Ggplot2: Elegant Graphics for Data Analysis. (Springer international publishing, Cham, 2016).

75. A. Lex, N. Gehlenborg, H. Strobelt, R. Vuillemot, & H. Pfister. UpSet: Visualization of Intersecting Sets. IEEE Trans. Vis. Comput. Graph. 20, 1983–1992 (2014).

76. Ritchie, M. E. et al. limma powers differential expression analyses for RNA-sequencing and microarray studies. Nucleic Acids Res. 43, e47–e47 (2015).

77. Kolde, R. pheatmap: Pretty Heatmaps. The R Foundation 10.32614/cran.package.pheatmap (2010).

78. Xu, S. et al. Using clusterProfiler to characterize multiomics data. Nat. Protoc. 19, 3292–3320 (2024).

79. Rappsilber, J., Ishihama, Y. & Mann, M. Stop and Go Extraction Tips for Matrix-Assisted Laser Desorption/Ionization, Nanoelectrospray, and LC/MS Sample Pretreatment in Proteomics. Anal. Chem. 75, 663–670 (2003).

80. Cox, J. & Mann, M. MaxQuant enables high peptide identification rates, individualized p.p.b.- range mass accuracies and proteome-wide protein quantification. Nat. Biotechnol. 26, 1367– 1372 (2008).

81. Tyanova, S. et al. The Perseus computational platform for comprehensive analysis of (prote)omics data. Nat. Methods 13, 731–740 (2016).

