## Supplementary Figures for "Structure-function dissection of huntingtin exon 1 identifies a PRD-driven modifier of neuronal toxicity in Huntington’s disease"

**This file includes:**

Extended Data Fig. 1 to 8

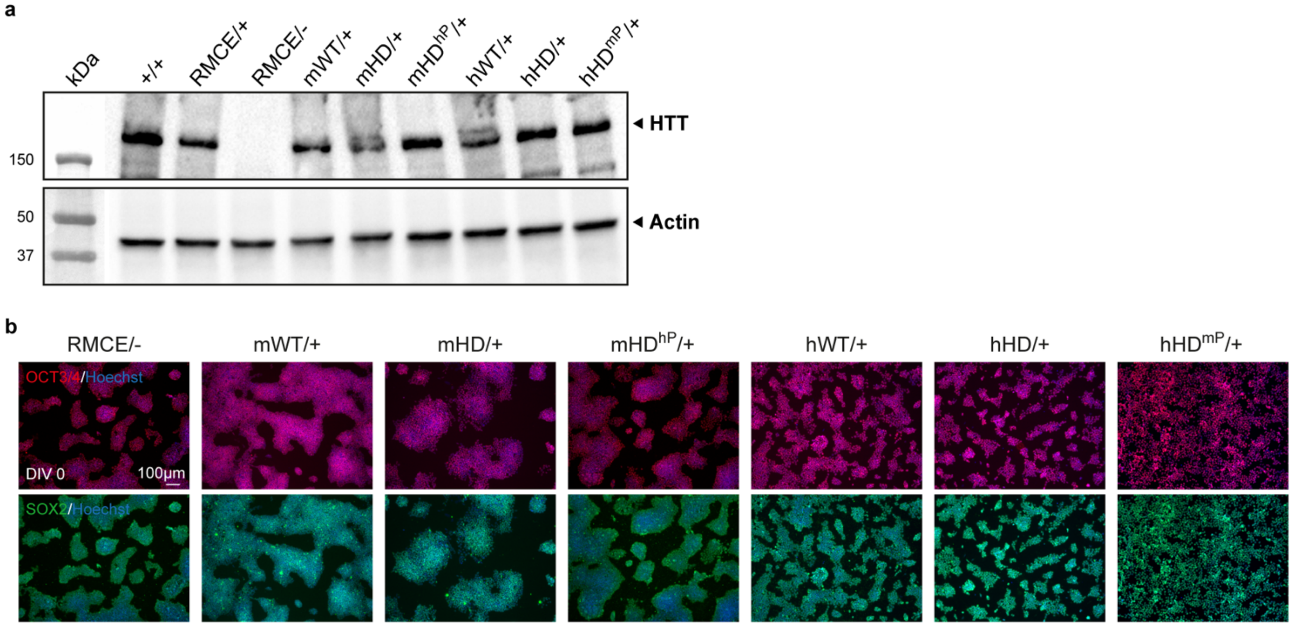


**Extended Data Fig. 1: Quality control on Htt knock-out and knock-in mESC lines**
**a**, Western blot analysis of huntingtin expression using the D7F7 antibody in E14 parental cells (+/+) and in Htt knock-out or knock-in mESC lines: RMCE/+, RMCE/-, mWT/+, mHD/+, mHD$^{hP}$/+, hWT/+, hHD/+ and hHD$^{mP}$/+ at the proliferative stage (DIV0). Actin was used as a loading control. **b**, Representative immunofluorescence images of RMCE/-, mWT/+, mHD/+, mHD$^{hP}$/+, hWT/+, hHD/+ and hHD$^{mP}$/+ mESC lines at DIV0, stained for OCT3/4 (red), SOX2 (green), and Hoechst (blue). Scale bar: 100 µm.

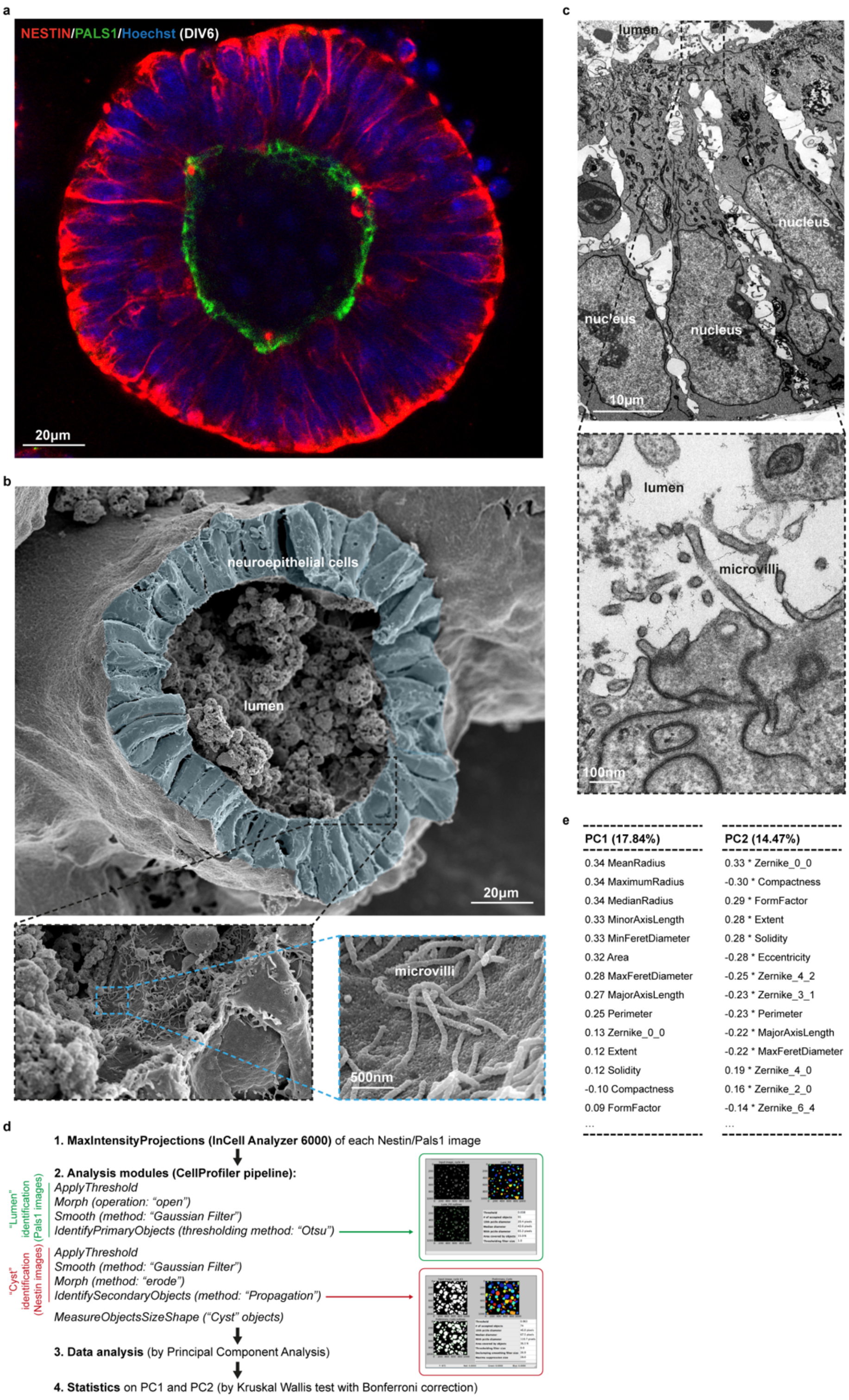


**Extended Data Fig. 2: Fluorescence and electron microscopy of neural cysts and detailed pipeline for automated multiparametric analysis**

**a**, Representative confocal image of a neural cyst derived from wild-type mESCs at DIV6, stained for NESTIN (red), PALS1 (green), and Hoechst (blue). The localization of the polarity marker PALS1 at the center of the neural cyst confirms correct apical-basal polarity. Scale bar: 20 µm. **b**, Scanning Electron Microscopy (SEM) image of a neural cyst at DIV7 derived from the E14 parental (+/+) mESCs. Scale bar: 20 µm. Zoomed inset: 500 nm. **c**, Transmission Electron Microscopy (TEM) of a neural cyst at DIV7 derived from +/+ cells. Scale bar: 10 µm. Zoomed inset: 100 nm. **d**, Schematic overview and detailed CellProfiler pipeline used for automated multiparametric analysis of neural cysts. **e**, Tables ranking individual neural cyst parameters by their contribution to PC1 and PC2, corresponding to the PCA shown in **Fig. 2d**.

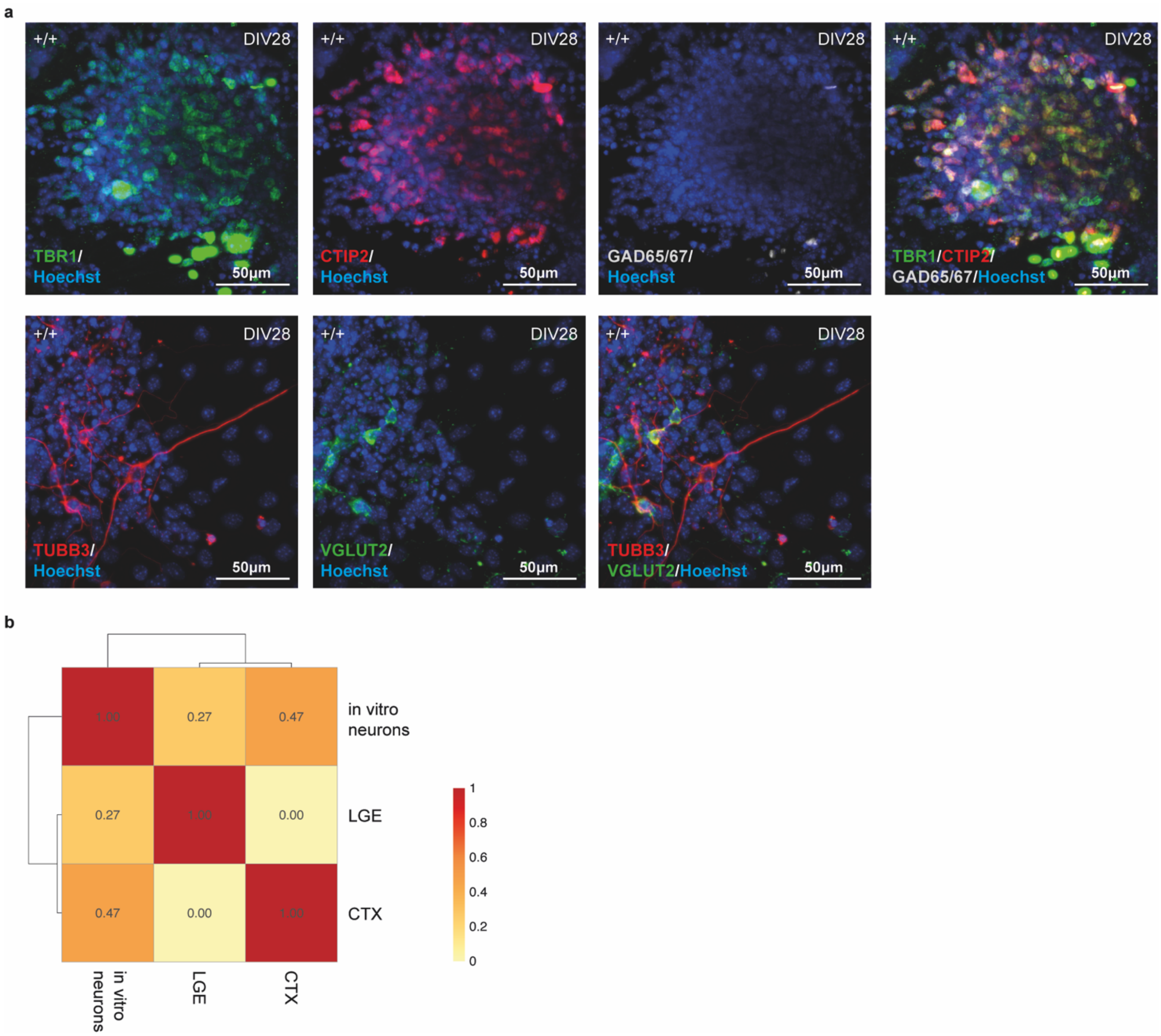


**Extended Data Fig. 3: Characterization of neuronal differentiation protocol**

**a**, Representative immunofluorescence images of wild-type (+/+) mESC-derived neurons at DIV29 of cortical differentiation. Top row: neurons stained for cortical markers TBR1 (green) and CTIP2 (red), the ventral marker GAD65/67 (white), and Hoechst (blue). Bottom row: neurons stained for the post-mitotic neuronal marker TUBB3 (red), the glutamatergic marker VGLUT2 (green), and Hoechst (blue). Scale bar: 50 µm. **b**, Heatmap of Jaccard similarity indices comparing the gene expression profile of the mWT/+ neuronal culture ("in vitro neurons") with reference gene sets from mouse fetal cortical ("CTX") or striatal (lateral ganglionic eminence, "LGE") tissues. The colour gradient indicates Jaccard similarity values (red: high similarity; yellow: low similarity).

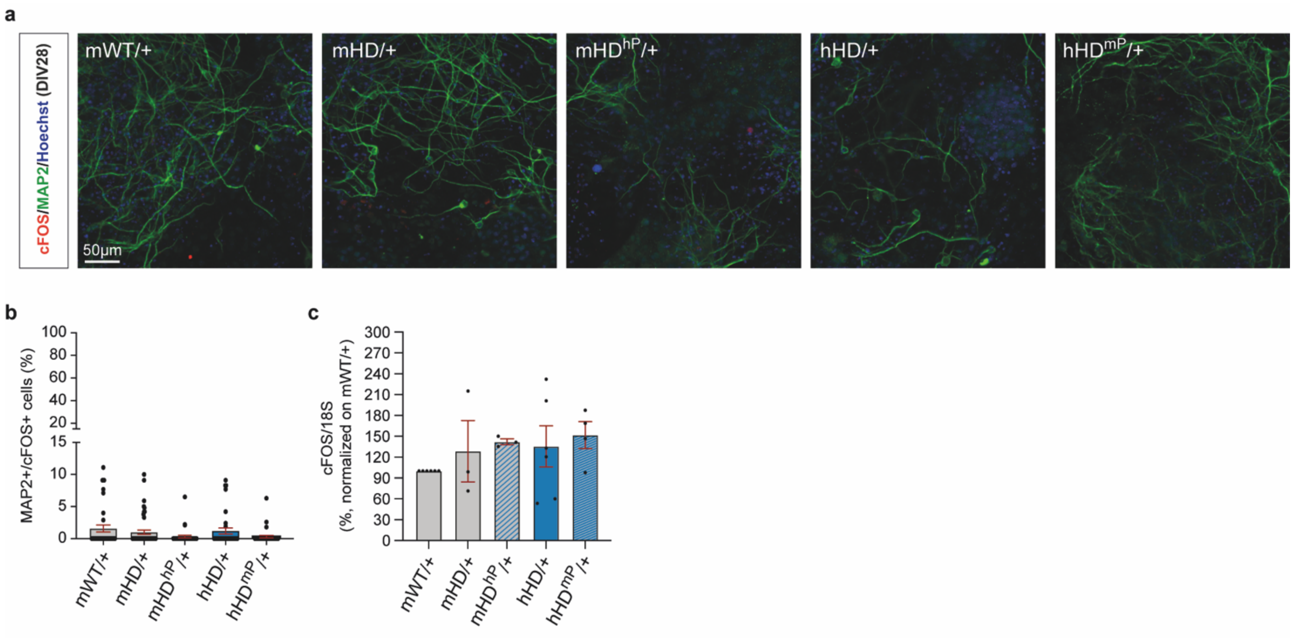


**Extended Data Fig. 4: Analysis of neuronal activity by cFOS quantification under basal conditions**
**a**, Representative immunofluorescence images of DIV28 neurons in basal condition (i.e. without glutamate stimulation), stained for cFOS (red), MAP2 (green), and Hoechst (blue). Scale bar: 50 µm. **b**, Quantification of immunofluorescence images showing the % of MAP2+/cFOS+ neurons per field. Each dot represents an individual field. Data are shown as mean ± SEM from n=3 independent experiments. **c**, qPCR analysis of cFOS expression in neuronal cultures under basal conditions, normalized to mWT/+. 18S was used as internal control. Each dot represents an independent qPCR replicate. Data are shown as mean ± SEM from n≥3 independent experiments.

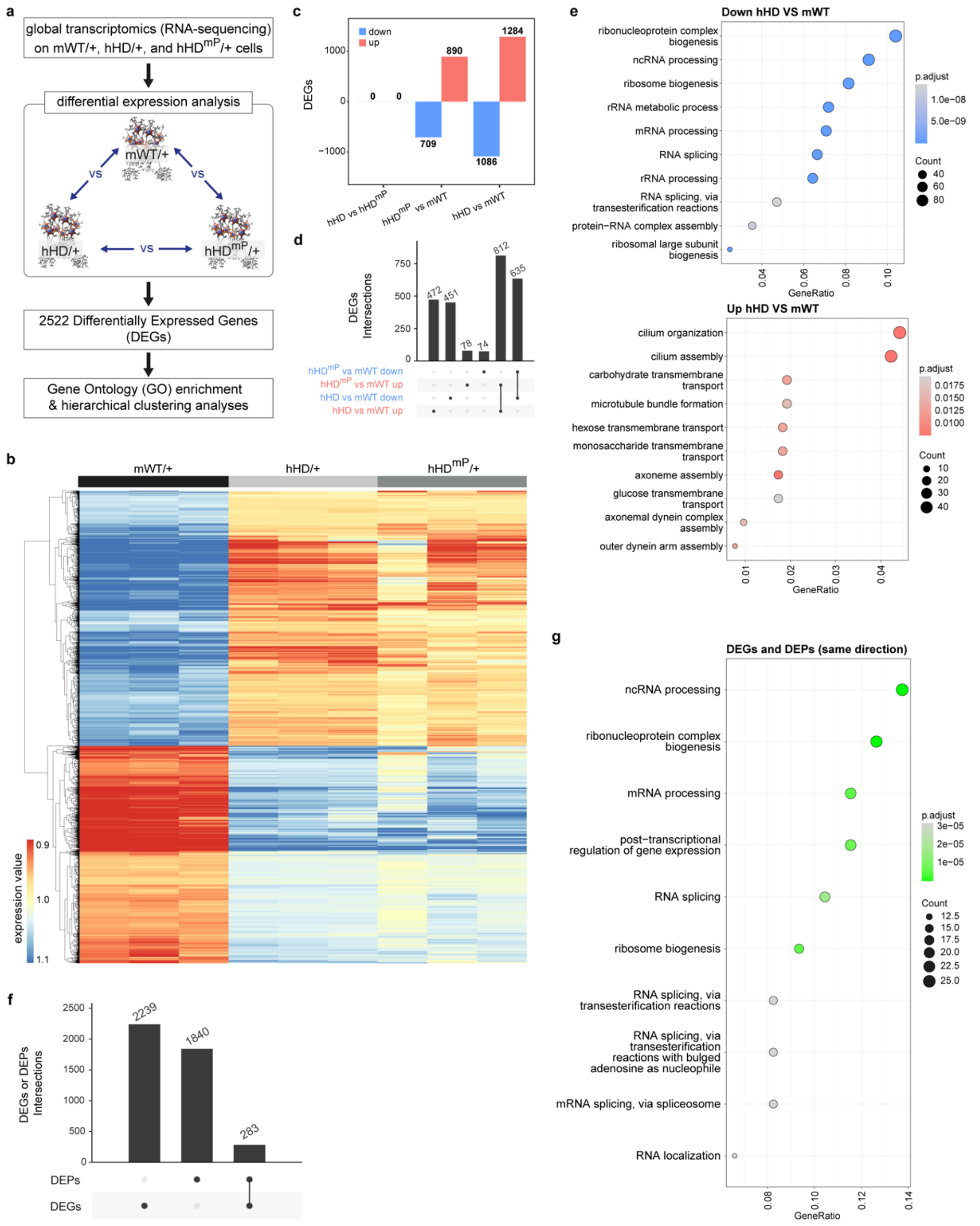


**Extended Data Fig. 5: Transcriptional profiling of mESC-derived neurons carrying different *HTT* exon 1 variants**

**a**, Schematic workflow of the transcriptomic analysis performed on neurons derived from mWT/+, hHD/+, and $hHD^{mP}$/+ mESC lines. **b**, Heatmap of hierarchical clustering of the 2,522 DEGs, showing normalized expression levels across genotypes. **c**, Bar plot showing the number of up- and downregulated genes in each pairwise comparisons. **d**, Upset diagram illustrating the intersections of upregulated (up) and downregulated (down) DEGs across different genotypes. Bars indicate the number of unique DEGs for each pairwise comparison (highlighted by a single dot below) or the number of overlapping DEGs between comparisons (highlighted by connected dots). **e**, Dot plots of enriched GO terms (ranked by gene ratio) for DEGs between mWT and hHD neurons. Dot size reflects the number of enriched genes; color represents the adjusted p-value. Up- and downregulated DEGs were analyzed separately. **f**, Upset diagram showing the intersection between DEGs and DEPs. Bars indicate unique DEGs or DEPs (single dots) and their overlap (connected dots). **g**, Dot plot of GO terms (ranked by gene ratio) enriched among DEGs/DEPs that are both differentially expressed and show consistent directionality between mWT and hHD neurons.

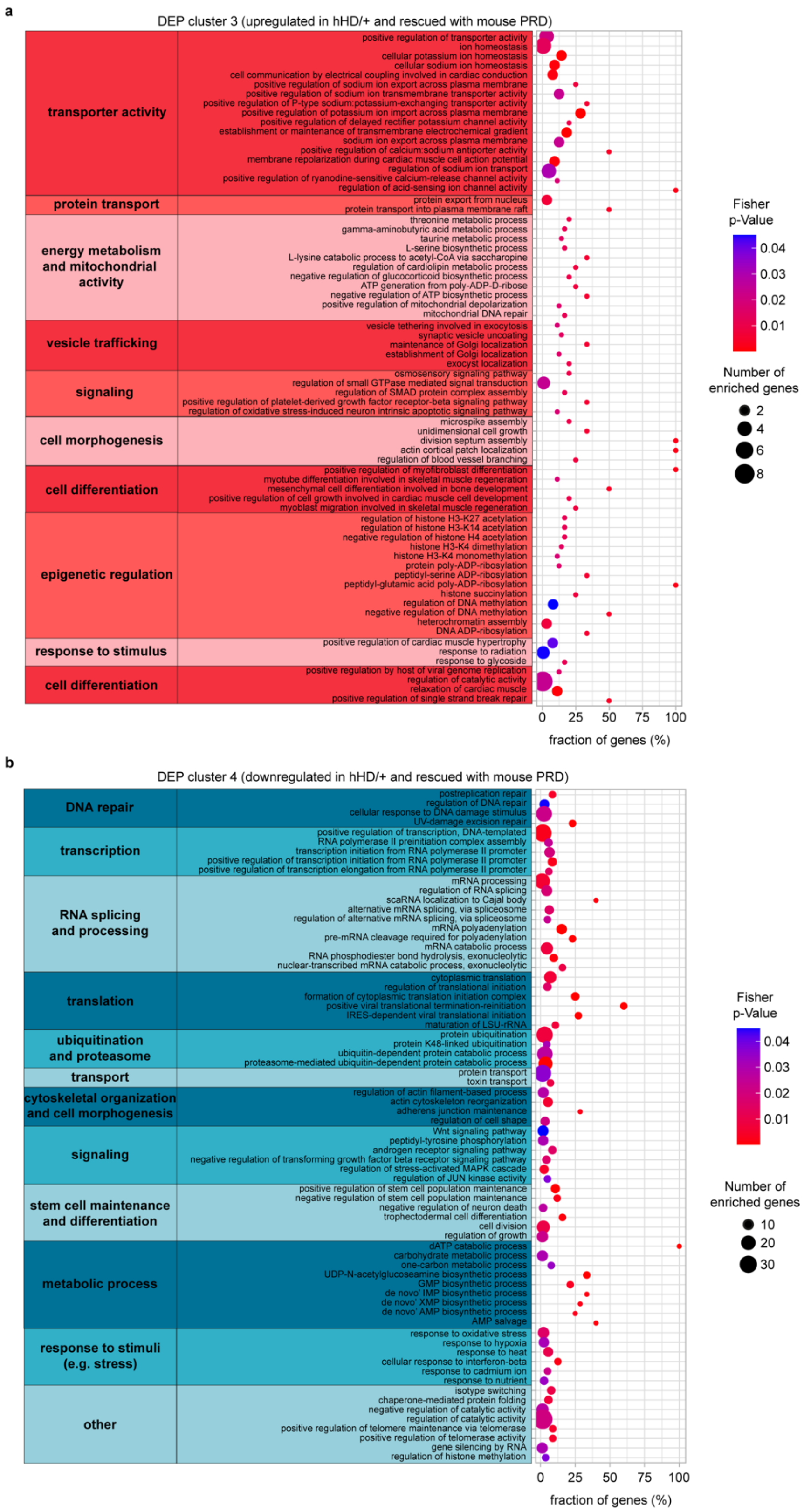


**Extended Data Fig. 6: GO enrichment analyses of rescued proteins**
**a**,**b**, Dot plots showing enriched GO terms (grouped into macro-categories) for DEPs that are either upregulated (**a**) or downregulated (**b**) in hHD/+ compared to the wild-type neurons, but whose expression is reverted in hHD$^{mP}$/+ neurons. These "rescued" proteins were identified in Clusters 3 and 4 of the clustering analysis shown in **Fig. 4b-d**. Each plot displays the gene ratio (fraction of enriched genes), number of enriched genes (dot size), and p-value (dot color) for each GO term. Upregulated and downregulated DEPs were analyzed separately.

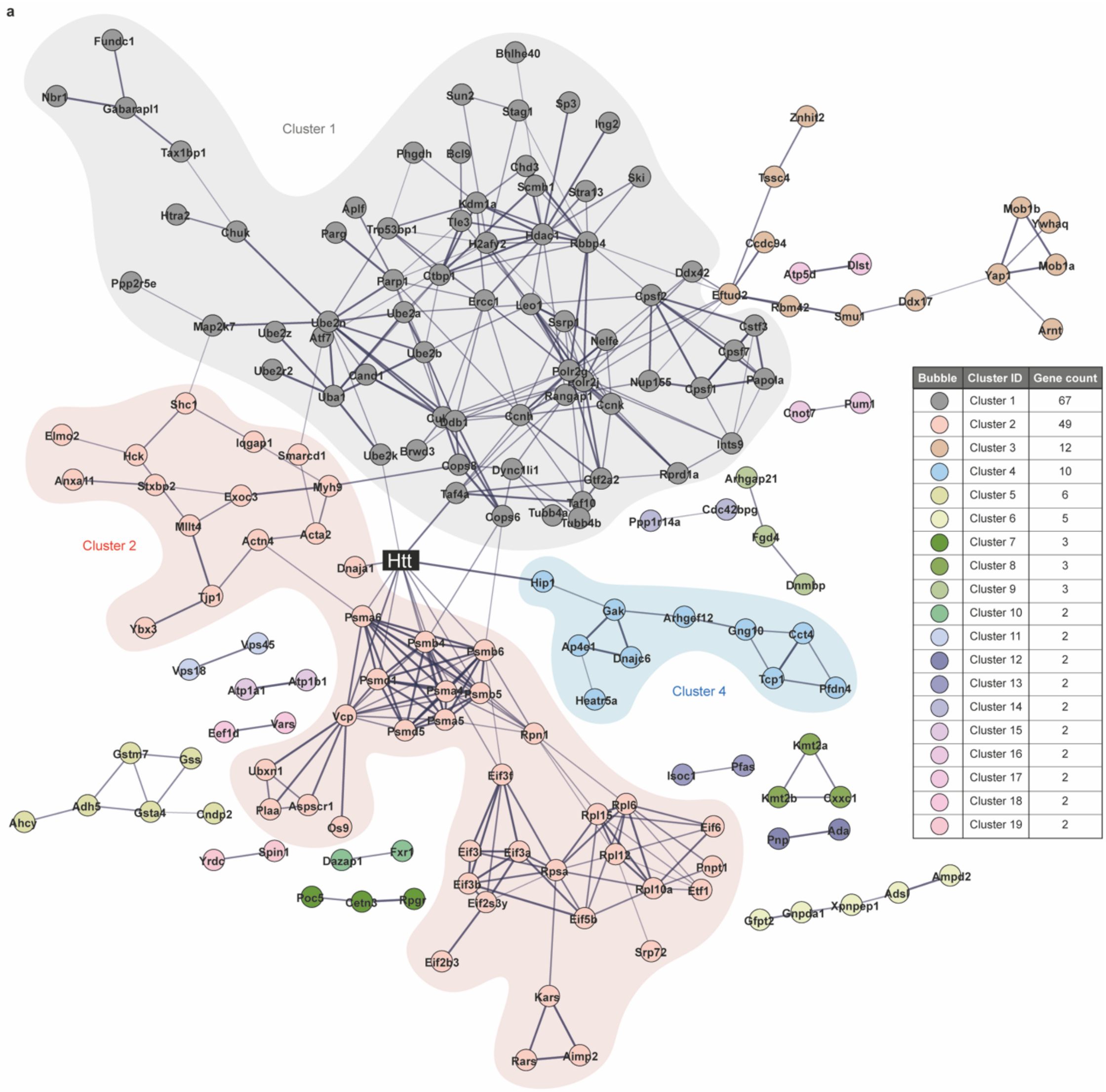


b

| Upstream regulator | Molecular type | n° of targets | Target molecules |
|---|---|---|---|
| HNF4A | transcription regulator | 51 | ACTA2;ACY1;ATF7;BLVRB;CCNH;CETN3;CHMP1B;CNN2;COPS8;CUL4B;CXXC1;DLST;DYNC1LI1;EFTUD2;FBLIM1;FKBPL;GOT1;GSS;HTRA2;INTS9;ISOC1;KARS1;MKRN1;NBR1;PAPOLA;PGM3;PLAA;PNP;PSMA5;PSMB5;PSMD1;RANGAP1;RBBP4;RBM42;RIOX2;RPL12;SAMHD1;SEPHS1;SKI;STOM;TARS1;TAX1BP1;TBCK;TLE3;TOM1;TPP2;TUBB4A;UBE2B;UBE2N;YJU2;ZNF277 |
| TP53 | transcription regulator | 48 | ACTA2;ACTN4;ADA;ADH5;AHCY;ATP1A1;BHLHE40;CAND1;CCNK;CD82;CHD3;CHUK;CNN2;CPSF1;DLST;DYSF;ERCC1;ING2;IPO9;KMT2A;MACROH2A2;MAP2K7;MYH9;PARP1;PGM3;PHGDH;PSMD1;RANGAP1;RBBP4;RPL12;RPN1;RPSA;SCMH1;SHC1;STAG1;STOM;THOP1;TIMM10;TJP1;TOM1;TUBB4A;TUBB4B;UBA1;UBE2B;UGDH;USP48;XPNPEP1;ZMAT3 |
| MYC | transcription regulator | 34 | ACTA2;ACTN4;AHCY;AIMP2;BZW2;CHKA;EFTUD2;EIF2B3;GDI2;GLG1;GOT1;MTHFR;PARP1;PFAS;POLR2G;PRDX4;PREP;RARS1;RBBP4;RIOX2;RPL10A;RPL12;RPL15;RPL6;RPSA;SP3;THOP1;TPP2;VARS1;VPS72;WWC1;YAP1;YBX3;YRDC |
| KRAS | enzyme | 27 | ACY1;ADGRL3;AIMP2;Akr1c14;ATP1B1;BZW2;EIF6;LLGL2;PALD1;PARG;PRDX4;QKI;QTRT1;RNH1;RPN1;RPSA;SAMHD1;SCMH1;STOM;TCP1;TIMM10;TLE3;TP53BP1;UBA1;VCP;XRN2;YAP1 |
| NFE2L2 | transcription regulator | 25 | ATF7;ATP1A1;CHORDC1;GOT1;GSS;Gsta4;NBR1;PHGDH;PRDX1;PREP;PSMA4;PSMA5;PSMA6;PSMB4;PSMB5;PSMB6;PSMD1;PSMD5;RARS1;SHC1;TP53BP1;UBE2K;UGDH;VCP;XRN2 |
| CTNNB1 | transcription regulator | 22 | ACTA2;ACTN4;Akr1c14;BCL9;BHLHE40;CNN2;DDX17;DNAJC6;FGD4;GFPT2;GNG10;IDI1;MFGE8;RPL10A;RPL12;RPL15;RPL6;RPSA;TJP1;UBA1;YAP1;ZNF277 |
| EGFR | kinase | 19 | ACTA2;ACY1;ADGRL3;BZW2;CHKA;EIF6;GNPDA1;IGF2BP3;MTHFR;PSMB4;PSMB5;PSMB6;QTRT1;RPN1;TIMM10;TUBB4A;TUBB4B;UBA1;XRN2 |
| APP | other | 19 | ACTA2;ATP1B1;ATP5F1D;CD82;CXXC1;GDI2;Gsta4;HARS1;HDAC1;HEATR5A;KMT2A;NBR1;RPGR;SHC1;SSRP1;TJP1;TOM1;TUBB4B;UBE2K |
| CD3 | complex | 15 | ACTN4;CCNH;ELAVL3;GDI2;ILF2;PFDN4;PNP;PRDX4;PREP;PSMA5;PSMA6;RARS1;RPL6;SHC1;STOM |
| HTT | transcription regulator | 15 | ACTA2;ATP1A1;BHLHE40;CHKA;CHUK;CPSF2;DLST;GSS;HTRA2;PSMA5;PSMB4;RABIF;SEPHS1;SP3;UBE2N |

**Extended Data Fig. 7: Interaction network and upstream regulators of rescued proteins**
**a**, Protein-protein interaction network map showing clustering of rescued proteins and their connections to Htt, based on STRING database analysis. Only experimentally validated and curated interactions with a confidence score >0.45 were included. Line thickness indicates the strength of experimental support. The table on the right lists all identified clusters and the number of proteins/genes in each. Clusters 1, 2, and 4 (highlighted in the network map) are directly connected to Htt. **b**, Summary table of upstream regulators identified by Ingenuity Pathway Analysis (IPA) using the QIAGEN database, performed on the set of rescued proteins. The top 10 regulators (out of 224 total) are shown, ranked by number of target molecules. The table includes each regulator's name, molecular type, number of targets, and associated downstream targets.

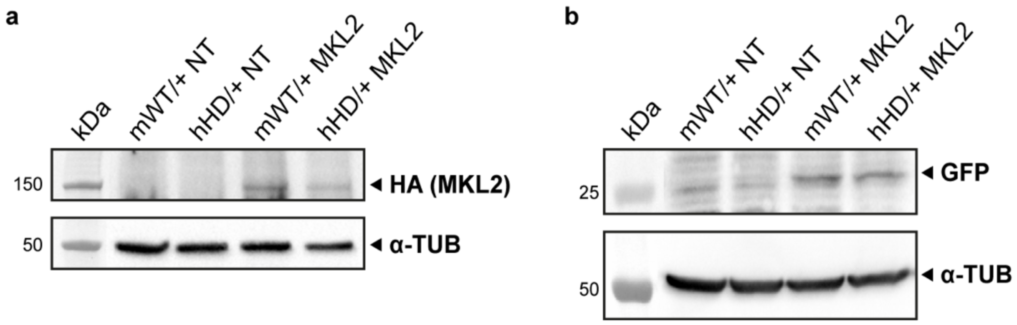


**Extended Data Fig. 8: Forced expression of MKL2 in maturing neurons**

**a**,**b**, Representative western blot analyses of HA-tagged MKL2 (**a**) and GFP (**b**) two days after plasmid transfection (at DIV14) in maturing neurons. α-Tubulin was used as loading control. NT: Non-transfected.
